# Role of histone variants H2BC1 and H2AZ.2 in H2AK119ub nucleosome organization and Polycomb gene silencing

**DOI:** 10.1101/2024.01.16.575234

**Authors:** Xiangyu Shen, Chunxu Chen, Yue Wang, Wan Zheng, Jiahuan Zheng, Amanda E. Jones, Bing Zhu, Hao Zhang, Charles Lyons, Arjun Rijal, James A. Moley, Gengsheng Cao, Kui Liu, Robert Winn, Amanda Dickinson, Kai Zhang, Hengbin Wang

## Abstract

Ubiquitination of histone H2A at lysine 119 residue (H2AK119ub) plays critical roles in a wide range of physiological processes, including Polycomb gene silencing ^1,2^, replication ^3–5^, DNA damage repair ^6–10^, *X* inactivation ^11,12^, and heterochromatin organization ^13,14^. However, the underlying mechanism and structural basis of H2AK119ub remains largely elusive. In this study, we report that H2AK119ub nucleosomes have a unique composition, containing histone variants H2BC1 and H2AZ.2, and importantly, this composition is required for H2AK119ub and Polycomb gene silencing. Using the UAB domain of RSF1, we purified H2AK119ub nucleosomes to a sufficient amount and purity. Mass spectrometry analyses revealed that H2AK119ub nucleosomes contain the histone variants H2BC1 and H2AZ.2. A cryo-EM study resolved the structure of native H2AK119ub nucleosomes to a 2.6A resolution, confirming H2BC1 in one subgroup of H2AK119ub nucleosomes. Tandem GST-UAB pulldown, Flag-H2AZ.2, and HA-H2BC1 immunoprecipitation revealed that H2AK119ub nucleosomes could be separated into distinct subgroups, suggesting their composition heterogeneity and potential dynamic organization. Knockout or knockdown of H2BC1 or H2AZ.2 reduced cellular H2AK119ub levels, establishing H2BC1 and H2AZ.2 as critical determinants of H2AK119ub. Furthermore, genomic binding profiles of H2BC1 and H2AZ.2 overlapped significantly with H2AK119ub binding, with the most significant overlapping in the gene body and intergenic regions. Finally, assays in developing embryos reveal an interaction of H2AZ.2, H2BC1, and RING1A *in vivo*. Thus, this study revealed, for the first time, that the H2AK119ub nucleosome has a unique composition, and this composition is required for H2AK119ub and Polycomb gene silencing.

## Main

Histone H2A lysine 119 ubiquitination (H2AK119ub) is the first discovered and most abundant ubiquitin conjugate in mammalian cells, occurring on 10-15% of cellular H2A ^15^. The functional significance of this modification was revealed by the identification of Polycomb Repressive Complex 1 (PRC1) as its ubiquitin ligase, linking this histone modification to Polycomb group (PcG) gene silencing ^1,11^. PcG proteins are a family of proteins that help maintain the repressed status of many master developmental regulatory genes that have wide-ranging roles in cell specification, epigenetic memory, genomic imprinting, and tumorigenesis ^16–20^. However, whether H2AK119ub is required for PcG gene silencing remains controversial. Studies in *Drosophila* revealed that point mutation of the ubiquitinating site in histone H2A gene did not impact PcG gene silencing; although mutant flies also exhibit lethal phenotype at a later embryonic stage ^21^. Consistently, PRC1 has been reported to compact chromatin physically and repress target gene expression, independent of its H2A ubiquitination activity ^22–27^. In contrast, accumulating evidence in mammalian cells underscores the essential roles of H2AK119ub in initiating the formation of Polycomb domains and PRC1-mediated gene repression ^28–33^. In *Drosophila*, H2AK119ub is distributed in transcription start sites (TSS), gene body, and intergenic regions ^34^. In mammalian cells, H2AK119ub has been mainly associated with the promoter regions of repressed genes ^31,32,35^. Low levels of H2AK119ub have also been observed throughout the genome ^35–37^. These genome-wide low levels of H2AK119ub are maintained by BAP1, which contributes indirectly to the high levels of H2AK119ub and PcG proteins at promoter regions ^36,37^. This explains the contradictory observation that BAP1 functions as a histone H2A deubiquitinase but it is required for PcG gene silencing ^38^. H2AK119ub has also been observed at active chromatin regions ^35,39–41^. The reasons underlying the divergent outcomes of H2AK119ub targets remain unclear.

Multiple models have been proposed to explain the repressive effects of H2AK119ub on gene expression. These models include antagonizing the active transcription histone marks H3K4me2/3 ^42^, excluding the binding of transcription elongation factor FACT ^43,44^, and recruiting linker histone H1 ^45,46^. H2AK119ub has been reported to recruit PRC2 to chromatin through its interactions with JARID2, an auxiliary subunit of Polycomb repressive complex 2 (PRC2) ^47,48^. JARID2 recognizes H2AK119ub *via* its ubiquitin-binding domain and recruits PRC2 to H2AK119ub-decorated regions. This process leads to the deposition of H3K27me3 and co-location of H3K27me3 and H2AK119ub marks ^29,47–49^. The deposited H3K27me3 then, through interacting with the chromodomain of the Cbx Polycomb subunit of canonical PRC1 (cRRC1) ^50^, recruits cRRC1 and organizes target regions into high-order chromatin structures ^23,25,51^. Mechanistically, PRC1 and H2AK119ub could repress target gene expression by controlling transcriptional burst frequency ^33^. Despite these advances, some important questions have been completely neglected from previous studies. For example, what is the composition of the H2AK119ub nucleosome? Is the composition important for H2AK119ub-mediated gene silencing? To answer these questions, we used the UAB domain of RSF1 to purify the H2AK119ub nucleosomes. Mass spectrometry analyses identified two histone variants, H2BC1 and H2AZ.2, in H2AK119ub nucleosomes. Subsequent studies revealed that H2BC1 and H2AZ.2 are indispensable for cellular H2AK119ub and PcG-mediated gene repression. Thus, this study revealed, for the first time, that the H2AK119ub nucleosome bears a unique composition, and this composition is required for the function of these specialized nucleosomes.

### Identification of histone variants H2BC1 and H2AZ.2 as components of H2AK119ub nucleosomes

In previous studies, we defined a small region in the remodeling and spacing factor 1 (RSF1) protein - the UAB domain (amino acids 770-807) - that interacts with the H2AK119ub nucleosomes specifically ^52^. To uncover the repressive mechanism associated with H2AK119ub, we purified mononucleosomes from 293T cells, using them as inputs in GST-UAB pulldown assays. GST-UAB, but not GST, pulled down H2AK119ub nucleosomes specifically, as evident by the depletion of H2AK119ub from Flowthrough and the enrichment of H2AK119ub in bound fractions (Figure 1a-1b, compare lanes 1-3 with 5-6). Parallel experiments with mononucleosomes purified from Ring1/Ring2 double knockout (DKO) 293T cells did not pull down any nucleosomes (Figure S1c-S1d, compare lane 5 with 6). Mass spectrometry analyses revealed that the major isoforms of histone H2A were K119 ubiquitinated at higher ratios in pulldown than in input nucleosomes (Figure 1c). Notably, H2AX, while being ubiquitinated in Input, did not interact with the UAB domain, and thus, no nucleosomes were pulled down (Figure 1c). Mass spectrometry analyses also identified two histone variants - H2BC1 (Q96A08) and H2AZ.2 (Q71U19) - that were highly enriched in GST-UAB pulldown H2AK119ub nucleosomes (Figure 1d). To confirm the mass spectrometry results, we established HA-Flag-H2BC1 and Flag-H2AZ.2 knockin (KI) mouse embryonic stem cell (mESC) lines with the CRISPR/Cas9 technology (Figure S2 and S3). Mononucleosomes were then isolated from these cells and used for the GST-UAB pulldown assay. When H2AK119ub was pulled down by GST-UAB, both Flag-H2BC1 and Flag-H2AZ.2 were also pulled down (Figure 1e and 1f, compare lane 3 with 2). Together, these experiments revealed that the H2AK119ub nucleosome has a unique composition, containing histone variants H2BC1 and H2AZ.2 as their components. Notably, not all H2BC1 and H2AZ.2 are associated with H2AK119ub. As shown in Figure 1e and 1f, when H2AK119ub was depleted from the Flowthrough, certain levels of H2BC1 and H2AZ.2 remained (panel B and C, compare lane 2 with 3).

**Figure 1.**
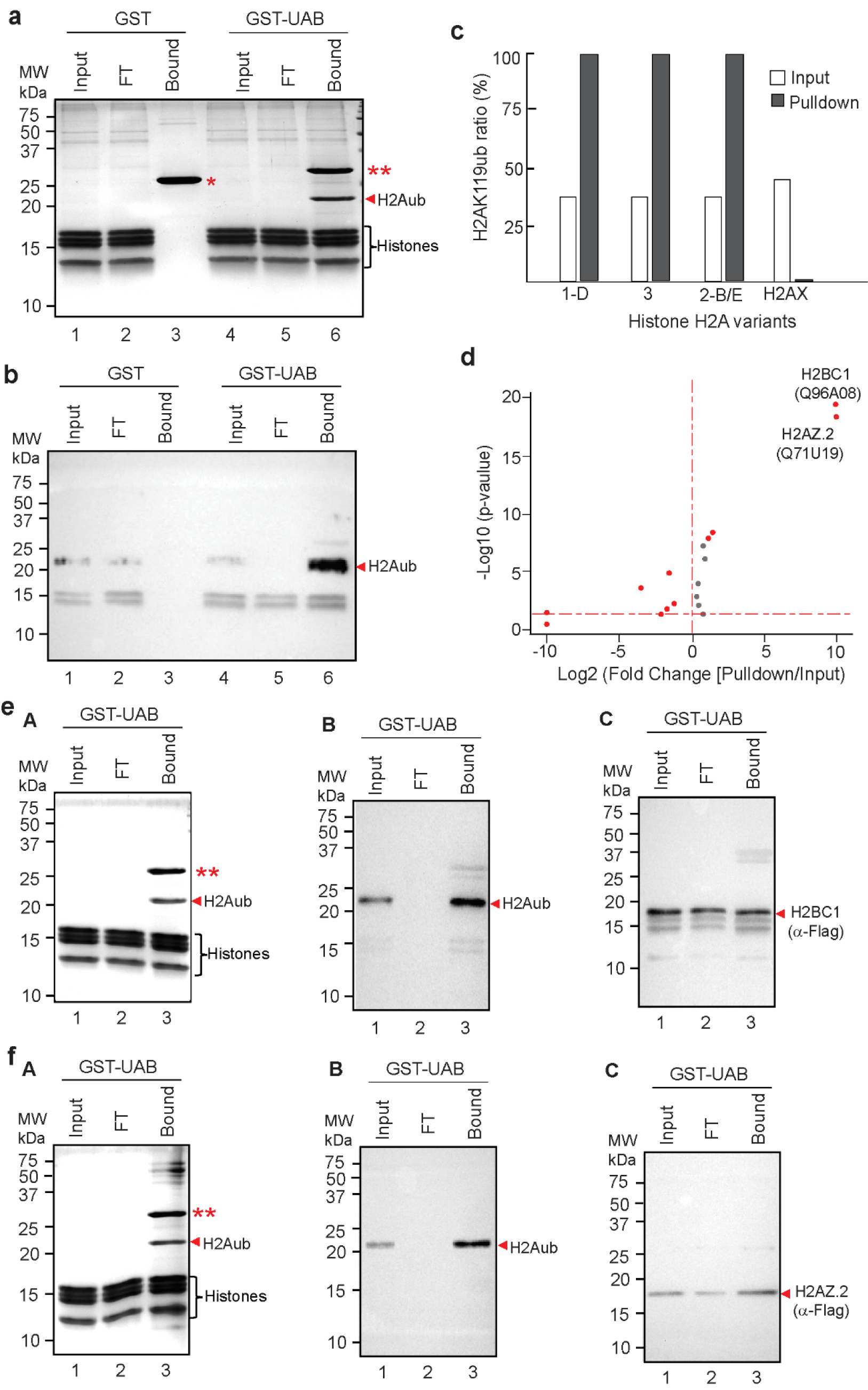
Identification of histone variants H2BC1 and H2AZ.2 as components of H2AK119ub nucleosomes. **a-b.** The UAB domain of RSF1 pulled down H2AK119ub nucleosomes specifically. Coomassie brilliant blue (CBB) staining (**a**) and anti-H2AK119ub immunoblots (**b**) of an SDS-PAGE containing an aliquot of Input, Flowthrough (FT), and Bound nucleosomes in GST (lane 1-3) and GST-UAB (lane 4-6) pulldown assays. H2AK119ub is indicated with red arrowhead. GST and GST-UAB are labeled with a red single and double asterisk, respectively. Molecular markers are shown on the left. The positions of core histones are labeled. **c.** Ratio of H2AK119 ubiquitination in Input and GST-UAB pulldown nucleosomes. The identified histone H2A variants are labeled. Notably, H2AX, although ubiquitinated in Input (50%), did not interact with the UAB domain, and no nucleosomes were present in pulldown samples. **d.** Volcano plot displaying mass spectrometry results of histone proteins enriched in pulldown nucleosomes from two independent experiments. *p* values calculated by two-sided *t*-test. **e-f.** CBB (**A**) and Immunoblots (**B, C**) of Input, FT, and bound mononucleosomes in an SDS-PAGE containing an aliquot of Input, Flowthrough (FT), and Bound nucleosomes in GST-UAB pulldown assays. Mononucleosomes were isolated from Flag-HA-H2BC1 (**e**) and Flag-H2AZ.2 (**f**) KI mESC. H2AK119ub is marked with a red arrowhead. GST-UAB is marked with red double asterisks. The positions of core histones are labeled.

### Structure of the native H2AK119ub nucleosome

To define the role of H2BC1 and H2AZ.2 in H2AK119ub nucleosome organization, we sought to determine the molecular interaction between these two histone variants with other histones within the H2AK119ub nucleosome. For this purpose, we first purified mononucleosomes from HeLa cells. The H2AK119ub nucleosomes were purified from total mononucleosomes by GST-UAB pulldown (Figure S4). Purified H2AK119ub mononucleosomes were cleaved from agarose beads by thrombin, further purified through a gel filtration Superdox 200 10/300 GL column, concentrated, and subjected to cryo-EM observation (Figure S4). The structure of native H2AK119ub nucleosome, precisely at 2.6 Å resolution, includes four histones H2A1B, H2BC1 (H2B1A), H3, H4, and a double-stranded DNA segment spanning 135 base pairs (Figure 2a-2c). Compared to reconstituted nucleosomes reported previously with X-ray (PDB:1AOI), the overall structure is strikingly similar. The 2.6Å cryo-EM structure, when superimposed with previously documented X-ray NCP structures (PDB:1AOI), displayed a root-mean-square deviation (RMSD) of less than 1Å (0.943), with closely aligned side chain locations. Within the nucleosome, H2A1B intimately envelops H2BC1 through multiple interacting regions (Figure 2dA). Tyr41 establishes a hydrogen bond with the oxygen in the DNA chain, while Asp69, in conjunction with Arg73, facilitates the linkage between H4 and H2A through hydrogen bonding interactions. The H2A1B L1 Loop engages in hydrogen interactions with H2BC1-T89 and S88, and these interactions are closely associated with nucleosomal DNA (Figure 2dB). The conserved acidic patch in our structure comprises histone H2A E56, E61, E64, D90, E91, E92, and H2B E114 (Figure 2dC). Since we used mononucleosomes for observation, interactions between nucleosome stacks, such as the contact point between H4 N-terminus and H2A-H2B acidic patch of adjacent nucleosomes, were not observed (Figure 2c). The interactions between the histones and DNA within the nucleosomes are intricately positioned (Figure 2c). H2A engages with DNA at the super helical location (SHL) 4 where the positively charged residues R11 and K13 intricately fit into the minor groove of the double-stranded DNA (Figure 2c, 2eA). Conversely, H2BC1 interacts with DNA at SHL3 where R34 engages with the minor groove, aided by K35, which forms contacts with the DNA backbone (Figure 2eB). Adjacent to SHL7, the DNA engages with histone H3, causing a distinct tilt away from the histone core (Figure 2eC). The H4 N-terminus, specifically R19, interacts with the DNA main chain of SHL2 (Figure 2eD). Our analyses of DNA sequences within H2AK119ub nucleosomes identified several conserved regions, which are particularly highlighted in the high stringency sequence in the SHL7 region (Figure 2f). Given the divergent C-terminus of H2A isoforms (Figure S5a) and the associated ubiquitin modification, we investigated the conformation of linker DNA bound to this region. We observed an extended H2A C-terminal tail (Figure 2g). Since recent cryo-EM studies revealed that nucleosomes reconstituted with H2A.Z.2 exhibit an open linker DNA conformation ^53^, our data suggest that H2A.Z.2 is absent in the subgroup of H2AK119ub nucleosomes whose structure we have resolved. The interaction between the Histone 2A C-terminal tail and H3 contributes to increased rigidity in this region, distinguishing it from the H2A.Z structure (PDB: 7M1X). Specifically, H3-Q55 forms hydrogen interactions with H2A’s C-terminal N110 while H3-I51 engages in hydrophobic interactions with H2A-I111. Additionally, H3-L48 and H2A-P117, L115, L116 create van der Waals forces and hydrophobic interactions (Figure 2g). The structure displays a distinct DNA-end on one side, contrasting with the opposite side that lacks DNA density, indicating the dynamic nature of the DNA terminus. Certain flexible regions, such as the C-terminal of H2A (119 to 127AA) and the N-terminus of H2B1A (1-32AA), remained elusive in our structure. As the ubiquitin modification occurs at the C-terminus of H2A, distanced from the core of the structure, the ubiquitin unit was also not observable.

**Figure 2.**
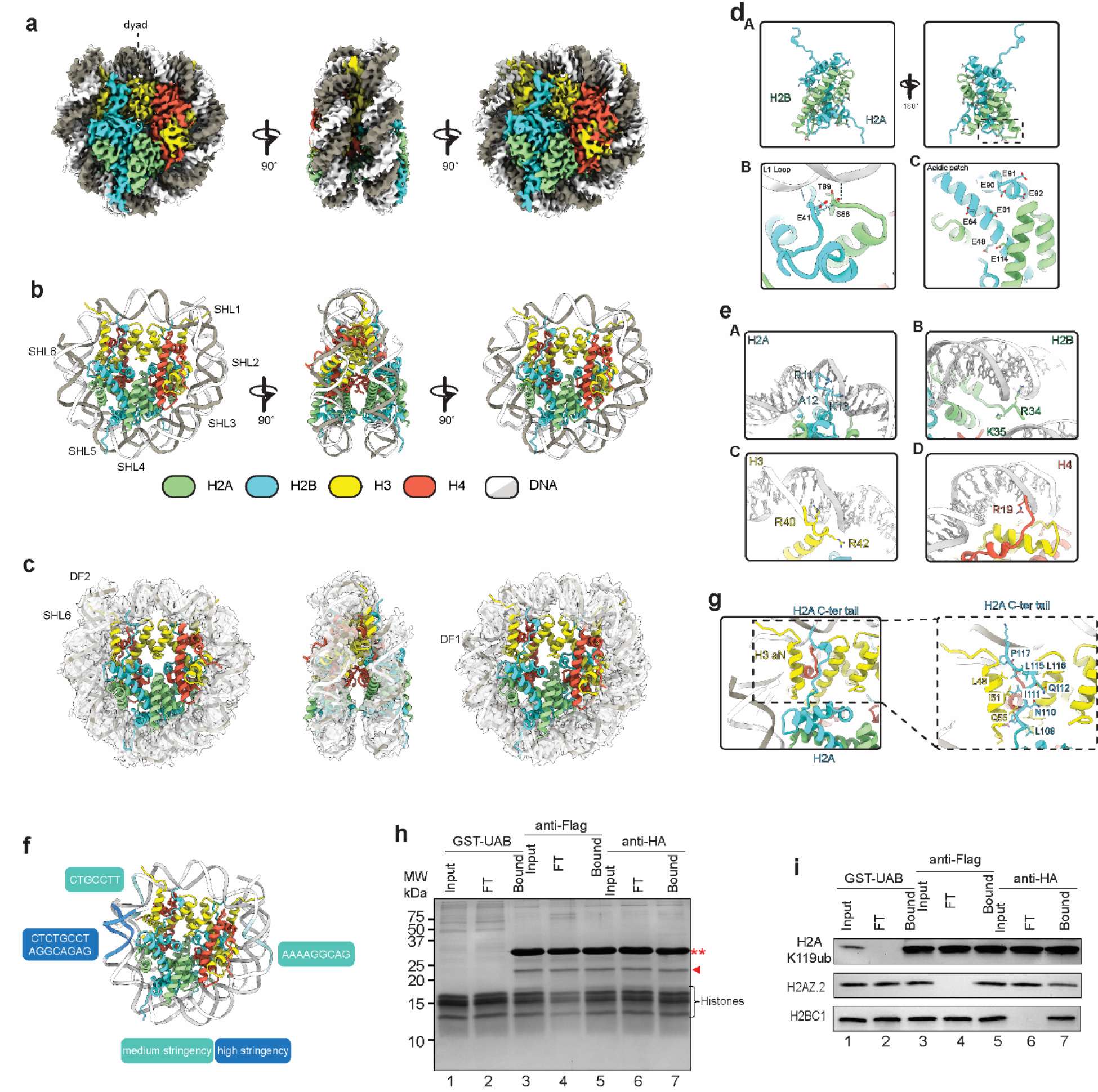
The structure of native H2AK119ub nucleosome. **a.** Cryo-EM density maps of H2AK119ub nucleosomes at 2.6Å, displaying disc view (left), side view (middle), and back view (right). **b.** Atomic model representation of H2AK119ub nucleosomes at 2.6Å, displaying disc view (left), side view (middle), and back view (right). **c.** Segmentation of the surrounding DNA oligomer into super helical locations (SHL) 0-7 and disc face (DL) 1-2. **d.** Interactions between H2A and H2B within H2AK119ub nucleosome structure. A. Comprehensive representation of multiple interactions between H2A and H2B, displayed from the disc view (left) and the back view (right). B. Detailed depiction of the specific interactions between H2A L1 loop and H2B. C. Identification and characterization of the conserved acid patch within our structure. **e.** Histone-DNA interaction in H2AK119ub nucleosomes. A. H2A engagement with DNA at SHL 4. B. H2B interaction with DNA at SHL3. C. DNA interaction with histone H3 adjacent to SHL7. D. Interaction between H4 N-terminus and the DNA main chain at SHL2. **f.** Identification of the high stringency sequence at SHL7 and medium stringency sequence within H2AK119ub nucleosomes. **g.** Enhanced rigidity resulting from the interaction between H2A C-terminus and H3. **h.** CBB staining of mononucleosomes purified from Flag-H2AZ.2 and HA-H2BC1 double tagged 293T cells subjected to sequential GST-UAB pulldown, anti-Flag and anti-HA IP. H2AK119ub is indicated with red arrowhead. GST-UAB is labeled with double asterisk. Molecular markers are shown on the left. The positions of core histones are labeled. **i.** Immunoblot of mononucleosomes purified from Flag-H2AZ.2 and HA-H2BC1 double tagged 293T cells subjected to sequential GST-UAB pulldown, anti-Flag and anti-HA IP. Protein identities are labeled on the left side of the panel.

Our biochemical studies identified two histone variants in H2AK119ub nucleosomes (Figure 1), yet our structural studies only identified H2BC1. We hypothesize that H2AK119ub nucleosomes are heterogenous, and there might be a dynamic organization and function diversification of these unique nucleosomes. To test this hypothesis, we have established a Flag-H2AZ.2/HA-H2BC1 dual tagged 293T cell line. When mononucleosomes purified from this cell line were subjected to sequential GST-UAB pulldown, anti-Flag IP, and anti-HA IP, we confirmed that H2AK119ub nucleosomes could be separated into subgroups containing H2BC1, H2AZ.2, and H2AZ.2/H2BC1 (Figure 2h-2i). Further work will need to reveal the structure and function of other groups of H2AK119ub nucleosomes. Nevertheless, our studies not only establish a robust experimental framework for future high-resolution studies of native nucleosomes but also corroborate our biochemical assays, which identified H2BC1 as a key component of the H2AK119ub-modified nucleosomes.

### H2BC1 and H2AZ.2 are required for cellular H2AK119ub

Since H2BC1 and H2AZ.2 are components of H2AK119ub nucleosomes, we investigated whether H2BC1 and H2AZ.2 are required for histone H2AK119 ubiquitination. For this purpose, we generated H2BC1 and H2AZ.2 knockout (KO) 293T cell lines using the CRISPR-Cas9 technology and confirmed the KO by sequencing of regions flanking the sgRNA target regions (Figure S6 and S7). As shown in Figure 3a and 3b, KO of either H2BC1 or H2AZ.2 abolished H2AK119ub. To further confirm this conclusion and determine whether the effects of H2BC1 and H2AZ.2 on H2AK119ub are conserved, we generated H2BC1 and H2AZ.2 KO mouse embryonic stem cell (ESC) lines with the CRISPR-Cas9 technology and confirmed the KO by sequencing of the genomic target regions (Figure S8 and S9). Similar to the situation in human 293T cells, KO of either H2BC1 or H2AZ.2 abolished H2AK119ub in mouse ESC (Figure 3c). Collectively, these data reveal that histone variants H2BC1 and H2AZ.2, as components of H2AK119ub nucleosomes, are required for cellular H2A K119 ubiquitination.

**Figure 3.**
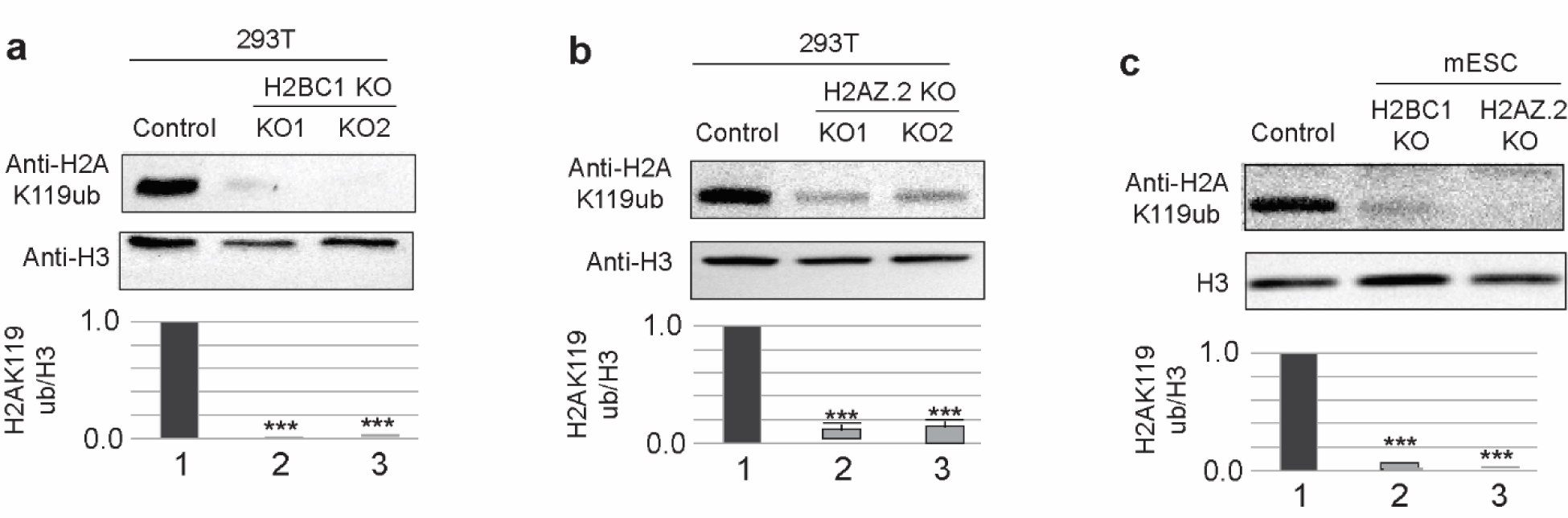
H2BC1 and H2AZ.2 are required for cellular H2AK119ub. **a.** H2BC1 is required for H2AK119ub in human 293T cells. Immunoblots of control and two independent H2BC1 KO 293T cell lines. Antibodies are labeled on the right side of the panel. Quantification of three biological replicates is shown at the bottom. Statistical differences were detected by one-way Anova. **b.** H2AZ.2 is required for H2AK119ub in human 293T cells. Immunoblots of control and two independent H2AZ.2 KO 293T cell lines. Antibodies are labeled on the right side of the panel. Quantification of three biological replicates is shown at the bottom. Statistical differences were detected by one-way Anova. **c.** H2BC1 and H2AZ.2 are required for H2AK119ub in mouse ESC. Immunoblots of control and H2BC1 and H2AZ.2 KO mouse ESC lines. Antibodies are labeled on the right side of the panel. Quantification of three biological replicates is shown at the bottom. Statistical differences were detected by one-way Anova.

### Genomic binding profiles of H2BC1 and H2AZ.2 overlapped significantly with H2AK119ub binding

Our biochemical studies revealed that, while H2BC1 and H2AZ.2 are components of H2AK119ub nucleosomes, not all H2BC1 and H2AZ.2 are associated with H2AK119ub (Figure 1e and 1f). To determine the extent of the genomic binding overlap between H2BC1 and H2AZ.2 with H2AK119ub, we measured the genome binding of H2BC1 and H2AZ.2, as well as H2AK119ub, using CUT&TAG technologies with the HA-Flag-H2BC1 and Flag-H2AZ.2 KI mouse ESC lines (Figure 1e and 1f). Data analyses revealed that H2BC1, H2AZ.2, and H2AK119ub are distributed widely in the genome, with more than half (57% for H2AK119ub, 58% for H2BC1, and 56% for H2AZ.2) located in gene or gene regulatory regions (promoter, 5’ UTR, exon, intron, 3’ UTR, TSS) and less than half (43% for H2AK119ub, 42% for H2BC1, and 44% for H2AZ.2) in intergenic and non-coding regions (Figure 4a). Notably, our data revealed that within the gene and gene regulatory regions, the major bindings are distributed in the intron and exon regions, with relatively minor fractions are distributed in the promoter and 5’UTR (H2AK119ub, 46% *vs* 9%; H2BC1, 47% *vs* 9%; H2AZ.2, 37% *vs* 18%) (Figure 4a). These data suggest that, in addition to the much-emphasized functions in promoters, H2AK119ub, and its associated H2BC1 and H2AZ.2, may play important regulatory roles in the gene body.

**Figure 4.**
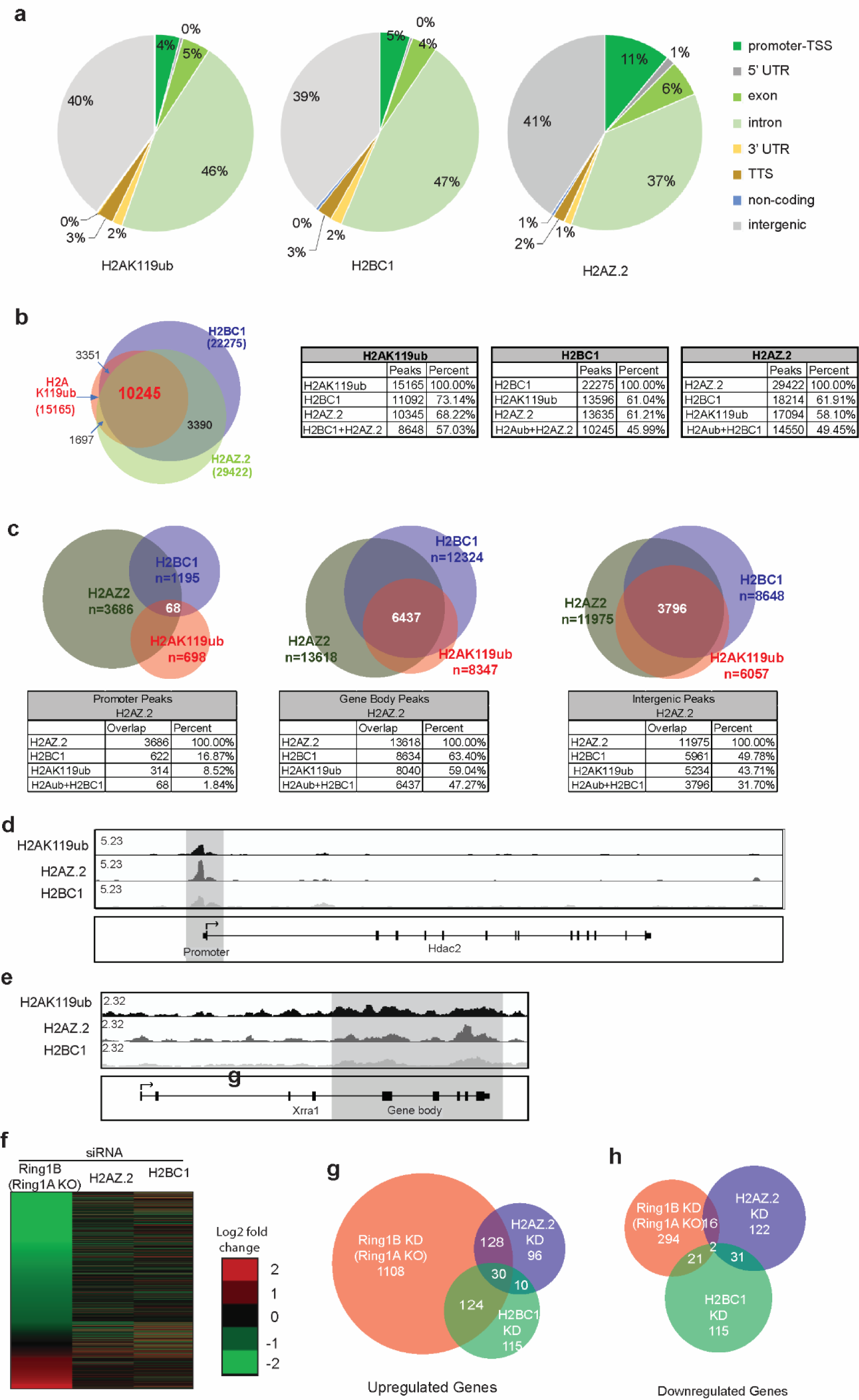
H2BC1 and H2AZ.2 genomic binding overlapped with H2AK119ub genome binding significantly. **a.** Pie charts showing the percentage of H2AK119ub (left), H2BC1 (middle), and H2AZ.2 (right) binding peaks to mouse reference genome (GRCh38.p12) following the RefSeq annotation assembly. The annotations are available from the genome database part of the NCBI database, and the data depicted above are from the Annotation Release 109. **b.** Left, Venn-diagrams showing overlaps of genome binding of H2AK119ub, H2BC1, and H2AZ.2. The numbers of peaks for each factor are labeled. Right, percentage of overlaps of genome binding peaks of H2AK119ub, H2BC1, and H2AZ.2. **c.** Top, Venn-diagrams showing overlaps of genome binding peaks of H2AK119ub, H2BC1, and H2AZ.2 at promoter region (left), gene body (middle), and intergenic regions (right). Bottom, percentage of overlaps of genome binding peaks of H2AK119ub, H2BC1, and H2AZ.2 at promoter region (left), gene body (middle), and intergenic regions (right) with H2AZ.2 as a viewpoint. **d.** Representative IGV track showing the overlap of H2AK119ub, H2BC1, and H2AZ.2 at the promoter of *Hdac2* gene. **e**. Representative IGV track showing the overlap of H2AK119ub, H2BC1, and H2AZ.2 at the gene body of *Xrra1* gene. **f.** Heat maps illustrating gene expression changes in mouse ESC in response to knockout/knockdown of Ring1A/Ring1B, H2BC1, and H2AZ.2. **g.** Venn-diagram showing overlaps of expression changes of up-regulated genes in mouse ESC in response to knockout/knockdown of Ring1A/Ring1B, H2BC1, and H2AZ.2. **h.** Venn-diagram showing overlaps of expression changes of down-regulated genes in mouse ESC in response to knockout/knockdown of Ring1A/Ring1B, H2BC1, and H2AZ.2.

Globally, the binding profiles of H2AK119ub overlap with H2BC1 and H2AZ.2 exclusively, with percentages of 68.22%, 73.14%, and 57.03% for H2BC1, H2A.2, and H2BC1/H2AZ.2, respectively (Figure 4b). Conversely, approximately 61.04% of H2BC1 overlapped with H2AK119ub, 61.21% with H2AZ.2, and 45.99% with both H2AK119ub and H2AZ.2 (Figure 4b). Similarly, about 58.10% of H2AZ.2 overlapped with H2AK119ub, 61.91% with H2BC1, and 49.45% with both H2AK119ub and H2BC1 (Figure 4b). These data correlate well with our biochemical studies and suggest that about half of H2BC1 and H2AZ.2 pools function independently of H2AK119ub (Figure 1). To pinpoint the specific regulation of H2AK119ub by H2BC1 and H2AZ.2, we analyzed the overlaps of the three factors in the gene promoter (promoter, TSS, 5’UTR), gene body (exon, intron, 3’ UTR), and intergenic region (non-coding and intergenic) (Figure 4c). Our results reveal that the overlapping ratios in gene bodies and intergenic regions are significantly higher than those in promoter regions, with 1.84% *vs* 47.27% and 31.7%, respectively (Figure 4c). Two representative IGV tracks of H2AK119ub, H2BC1, and H2AZ.2 overlaps at promoter region and gene body were shown in Figure 4d and 4e, respectively. Together, these data suggest that H2AK119ub nucleosomes may have additional functions beyond gene promoters.

Since H2BC1 and H2AZ.2 regulate H2AK119ub, we determined whether H2BC1 and H2AZ.2 regulate genes similar to those regulated by H2AK119ub. For this purpose, we measured gene expression changes in response to H2BC1 KD and H2AZ.2 KD in mouse ESC and compared to that of Ring1A KO/Ring1B KD. RNA-seq results revealed that while upregulated genes were affected more than down-regulated genes by H2BC1, H2AZ.1, and Ring1A/Ring1B, consistent with the gene repressive function of H2AK119ub (Figure 4f-4h), the overlap ratios are relatively low. Moreover, the high levels of overlap within the gene body regions suggest that H2AK119ub, together with its associated H2BC1 and H2AZ.2, may regulate other aspects of chromatin function. When analyzing the splicing efficiency of genes bound by H2AK119ub, H2BC1 and H2AZ.2, Ring1AKO/Ring1BKD, H2BC1 KD and H2AZ.2 KD resulted in significantly decreased splicing efficiency only when bound to the gene bodies of the respective genes (Figure S10). Thus, these studies pointed out a new function of H2AK119ub in regulating the splicing efficiency when bound to gene body.

### Function interaction of H2AZ.2, H2BC11 and RING1A in developing *Xenopus* embryos

To investigate whether the H2BC1 and H2AZ.2 regulation of H2AK119ub is conserved and functions *in vivo*, experiments were performed in the developmental model *Xenopus laevis*. First, *Xenopus* H2BC1 and H2AZ.2 homologs were identified. Using the *Xenopus* cell line XL-58, we purified *Xenopus* H2AK119ub nucleosomes by GST-UAB pulldown (Figure 5a). Similar to the situation in mammalian cells, GST-UAB specifically enriched H2AK119ub from input nucleosomes. Mass spec analyses identified two variants, H2BC11 (also called H2B1.1) and H2AZ.2, in H2AK119ub nucleosomes (Figure 5a). Evidence that these histone variants share a conserved function with their human counterparts is the high degree of sequence similarity, *Xenopus* H2AZ.2 shares 100% identity with the human protein while *Xenopus* H2BC11 protein shares a 92% identity to human H2BC1. Supporting a connection between H2AZ.2, H2BC11 and the PRC1-H2AK119ub pathway is data showing that these proteins together with RING1A are expressed at markedly similar dynamics over early development in *Xenopus* embryos (Figure 5b). If H2AZ.2, H2BC11, and RING1A interact during development, then we hypothesized that knockdown of all three proteins together would synergize and cause malformations. To test this hypothesis, a sensitized phenotypic assay was performed where low sub-phenotypic concentrations of each H2AZ.2, H2BC11, and RING1A MOs were injected into embryos at the one-cell stage. The majority of embryos injected with low concentrations of H2AZ.2 MOs or H2BC11 MOs or RING1A MOs alone appeared similar to embryos injected with control MOs (compare Figure 5ci-iv). However, when H2AZ.2, H2BC11, RING1A MOs were combined, keeping the total volume and concentration of MOs the same as controls, the majority of these triple morphants died and those that survived were abnormal with a short, curved axis (Figure 5cv-vi). Together, these results provide evidence for a functional role of H2BC11/H2AZ.2 in regulating H2AK119ub-mediated development in *Xenopus* embryos.

**Figure 5:**
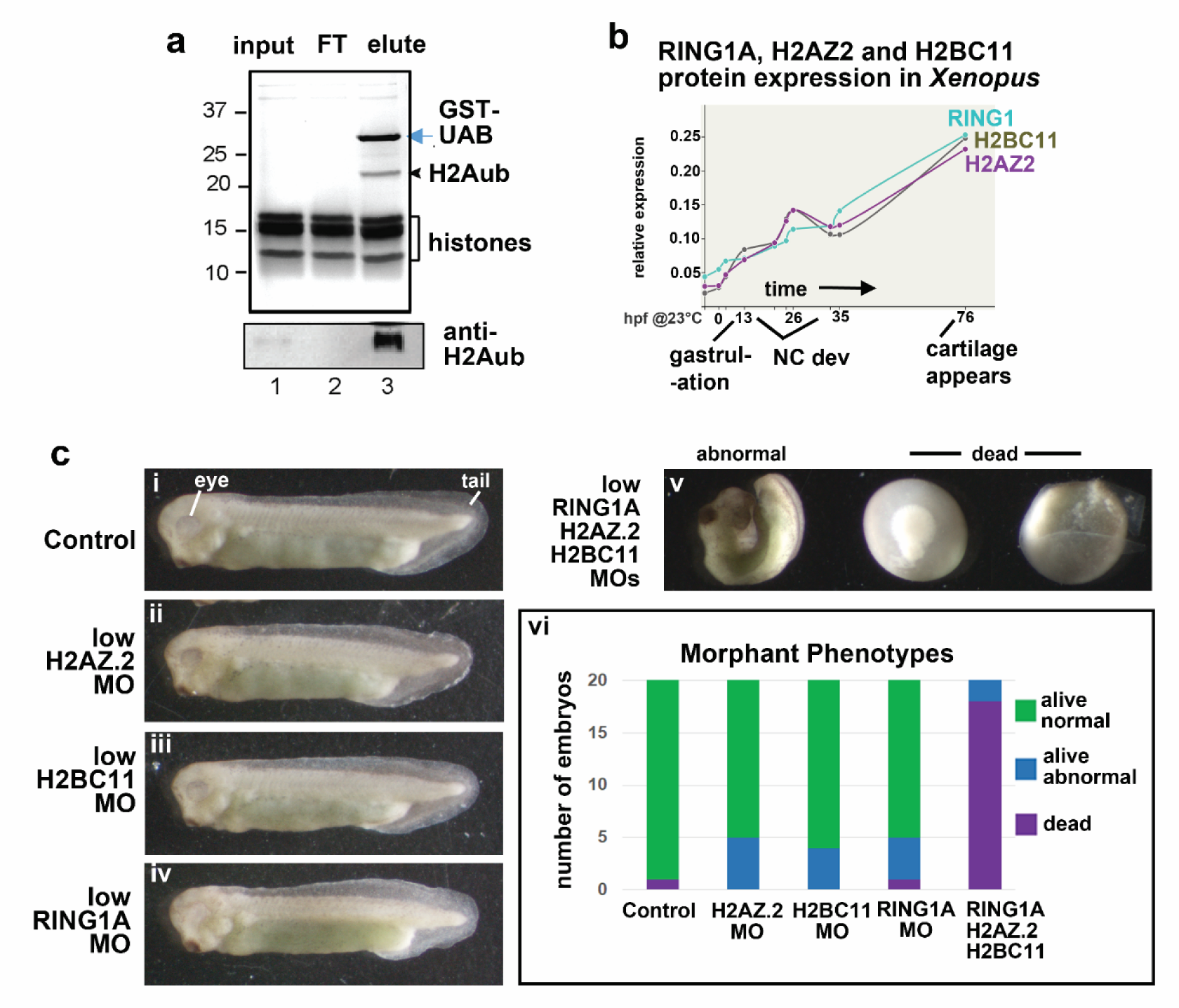
Function interaction of H2BC11, H2AZ.2, and RING1A in developing *Xenopus* embryos. **a.** Purification of *Xenopus* H2AK119ub nucleosomes by GST-UAB pulldown from the cell line XL-58. **b.** Relative protein expression of H2BC11, H2AZ.2, and RING1A over early development up to 76 hours post fertilization (hpf) at 23°C. NC= neural crest. Data generated on Xenbase.org ^87^. **c.** Lateral views (anterior to the left) of representative embryos injected with MOs to test for functional interaction. i) Embryo injected with 30ng/embryo if standard control MO. ii) Embryo injected with 10ng/embryo if H2AZ.2 MO and 20ng/embryo of control MO. iii) Embryo injected with 10ng/embryo H2BC11 and 20ng/embryo of control MO. iv) Embryo injected with 10ng/embryo RING1A and 20ng/embryo of control MO. v) Three representative embryos injected with a combination of 10ng/embryo of each H2AZ.2 MOs, H2BC11 MOs, RING1A MOs (30ng/embryo total). The first embryo has an axis defect and the second two embryos are dead. vi) Embryos from each injection group were divided into 1) alive and normal, 2) alive and abnormal and 3) Dead and then visualized in a stacked bar graph (n=20 embryos).

## Conclusion

The incorporation of histone variants, the nonallelic isoforms of canonical histones, alters nucleosome properties and endows auxiliary epigenetic information ^54^. Until this study, it was assumed that H2AK119ub-modified nucleosomes have the same subunit composition as other nucleosomes. Consequently, canonical histones, together with semi-synthetic or synthetic H2AK119ub proteins, were used to reconstitute nucleosomes and characterize its biochemical and biophysical properties ^44,55,56^. In this study, we took advantage of the H2AK119ub-specific binding (UAB) domain of RSF1 to purify H2AK119ub nucleosomes, and revealed, for the first time, that H2AK119ub nucleosomes have a unique composition, containing two histone variants H2BC1 and H2AZ.2. Importantly, these histone variants are required for cellular H2AK119ub. Although histone modifications have long been recognized as an important mechanism for PcG protein-mediated gene silencing, the role of specific histone variants in PcG activity has not been investigated before. H2BC1 (alternative name: H2B1A, TH2B) is highly expressed in early spermatocytes and replaces 80% of the major H2B prior to histone-protamine transition required for spermiogenesis ^57^. H2BC1 is also highly expressed in oocytes, and when over-expressed together with testis-specific histone H2A variant and phosphorylated nucleophosmin 1, it can induce open chromatin conformation and enhance induced pluripotent stem cell generation ^58,59^. H2AZ proteins have previously been implicated in PcG silencing, but the isoform and mechanisms were not identified ^60^. The identification of H2BC1 and H2AZ.2 as H2AK119ub nucleosome components not only reveals novel and critical roles for these histone variants but also uncovers nucleosome compositions as an important mechanism for Polycomb gene silencing.

H2BC1 and H2AZ.2 may confer special features to H2AK119ub nucleosomes. H2BC1 has been shown to alter interactions between histones ^61,62^; thus, the incorporation of H2BC1 in H2AK119ub nucleosome may facilitate nucleosome structural transition. While we did not observe H2AZ.2 in H2AK119ub nucleosome structure, H2AZ.2 nucleosomes have been shown to more stable than H2AZ.1 nucleosome, despite the fact that these isoforms only differ by three amino acids ^53^. It should be noted that all previous studies used recombinant histones and strong nucleosome positioning sequences. These studies revealed a highly conserved and relatively stable structural fold that appears insensitive to histone variant incorporations. In contrast, this study, for the first time, solved the high-resolution structure of native nucleosomes. The difference between the structure of native and reconstituted nucleosomes may come from both its histone compositions, but also the DNA used in previous reconstitution experiments. These DNA used in nucleosome reconstitution often contains strong nucleosome positioning sequences and may cause diverged results from the native nucleosome formation. The relative homogeneity of H2AK119ub nucleosome has allowed us to resolve the native nucleosome to high resolution. We envision that proteins or domain that bind specifically to a particular subgroup of nucleosomes may help overcome the heterogeneity associated with native nucleosomes and open a new era of nucleosome structural study, revealing the *bone fide* connection between nucleosome structure (composition) and function.

Previous studies on H2AK119ub in mammalian cells primarily focused on its repressive function at promoter regions. However, our current research reveals that in additional to promoter, H2AK119ub may also function in gene coding regions, including exons and introns. Our study revealed that Ring1AKO/Ring1BKD, H2BC1 KD and H2AZ.2 KD resulted in decreased splicing efficiency only when bound to the gene body of the respective genes. Since the difference is not dramatic, we reason that these could be indirect effects of Pol II processivity.

Nonetheless, the precise function of H2AK119ub in gene coding regions remains to be elucidated. Moreover, H2AK119ub in gene coding regions exhibits a higher overlap ratio with H2BC1 and H2AZ.2 compared with H2AK119ub in promoter regions. These findings suggest potential distinct functions of H2AK119ub nucleosomes in promoter *vs* coding regions. In addition to its role in gene regulation, H2AK119ub is widely distributed in intergenic regions. While the function of H2AK119ub in these regions remains elusive, its presence has been observed in *Drosophila* cells ^34^. Insights may be drawn from the studies of mammalian *X* inactivation ^63^. H2AK119ub first accumulates in intergenic regions on the inactivated *X* chromosome, and then spreads to gene bodies, promoters, and putative enhancers, during *X* inactivation. The patterns observed during *X* inactivation raise questions about whether H2AK119ub in autosomes follows similar patterns and whether H2AK119ub at intergenic regions plays a role in regulating genomic stability or higher-order chromatin structure. Elucidating the function and mechanism of H2AK119ub in intergenic regions will be future tasks in PcG studies and the epigenetic field.

## Methods

### Cells and cell culture

Human Embryonic Kidney 293T cells were purchased from ATCC (CRL-3216, Manassas, VA, US). HeLa S3 cells were described previously ^1^. 293T and Hela cells were cultured in DMEM medium (Corning, NY) supplemented with 10% FBS (FCS, R&D Systems, S11550) and 1% Ampicillin-Streptomycin (Gibco, Invitrogen). Mouse embryonic stem cell line R1 was purchased from ATCC (CB-SRRC-1011) and cultured in ESC media, containing DMEM medium (Corning, NY, MT010-013-CM) supplemented with 50 unit per ml penicillin and streptomycin (Gibco, Invitrogen), 15% murine ESC defined FBS (R&D System, S10250), 1% GlutaMax (Gibco, Invitrogen), 1 X nonessential amino acids (Corning, NY, MT-25-025-CI), 1 X nucleoside (Millipore, ES-008-D), 0.007% b-mercaptoethanol (Sigma, M3148), and 1,000 unit/ml mLIF (Millipore, ESGRO) on 0.1% gelatin coated plates or irradiated mouse embryonic fibroblasts (Millipore, PMEF-NL) and passaged every other day as described previously ^64^.

To establish Flag-H2AZ.2/HA-H2BC1 double-tagged 293T cell lines, human H2BC1 and H2AZ2 cDNA were synthesized by Thermo Fisher and cloned into the pCW-Cas9 vector (Addgene #50661) by replacing the Cas9 gene. Lentiviruses were packaged as follows: 7 µg of the plasmids encoding Flag-H2AZ.2 or HA-H2BC1, 3 µg of psPAX2 (Addgene #12260), and 1.5 µg of the pMD2.G plasmid (Addgene #12259) were mixed with 13 µl Lipofectamine 3000 and 13 µl P3000 (Invitrogen) in 600 µl of Opti-MEM reduced serum medium (Thermo Fisher). After incubation at room temperature for 15 minutes, the mixture was added to HEK293T cells in a 10 cm cell plate containing 8 ml of DMEM medium with 10% fetal bovine serum. Cell culture medium was harvested at 24 and 48 hours respectively, centrifuged at 1000g for 10 minutes to remove cell debris, and filtered through a 0.45 µm filter. Lentivirus in the culture medium was concentrated using a lentivirus concentrator (Clontech Labs 3P #631231). One-third of the concentrated virus was used to infect 2.5 x 10^5^ 293T cells in the presence of 0.8 mg/ml polybrene. After 2 days of infection, cells were selected with puromycin for 6 days and then diluted into 96-well plates to obtain single clone. Individual clones were expanded and analyzed by immunoblots with antibodies against the epitope tags. Tet-on Flag-H2AZ2 stable cells were established first, and double-tagged cells were obtained by transducing Tet-on Flag-H2AZ2 stable cells with lentivirus encoding Tet-on HA-H2BC1.

### Nucleosome isolation, GST-pulldown assay, and cryo-EM sample preparation

To purify mononucleosomes, nuclei from 2 X 10^8^ Hela or 293T cells were digested with 20 U of micrococcal nuclease (Worthington Biochemical Corporation, LS004797) for 45 minutes, and nucleosomes were extracted following a previously described protocol ^65^. Extracted nucleosomes were diluted with histone storage buffer (10 mM Hepes-KOH pH 7.5, 1 mM EDTA, 60 mM KCl, 10% glycerol, 0.2 mM PMSF) to a final concentration of 0.35 mg/ml and dialyzed against histone storage buffer overnight.

For GST-UAB protein purification, the sequence of the UAB domain (amino acids 770-807 of RSF1 protein) were cloned into the pGEX-4T1 (ATCC, 77103). GST and GST-UAB were purified from BL-21 cells with GST agarose (Thermo Scientific, PI20211) following the manufacture’s instruction. Two micrograms of GST or GST-UAB were bound to 50 microliter GST-agarose in 1 x PBS buffer. After washing with histone storage buffer, the GST or GST-UAB-bound agaroses were incubated with mononucleosomes at 4°C for 1 hr. The GST-agarose beads were then washed with histone storage buffer for 3 times, and the bound proteins were eluted with reduced glutathione, and used for CBB, immunoblots, or mass spectrometry assays. DNAs associated with H2AK119ub nucleosomes and input nucleosomes were extracted, and resolved on 1.2% agarose gel ^65^. Gel bands corresponding to mononucleosomes (100-200 bp) were excised and DNAs were purified and subjected to deep-sequencing ^65^.

For Cryo-EM sample preparation, the extracted nucleosomes were separated on a 5-30% linear sucrose gradient. The protein and DNA profiles of sucrose gradient were analyzed with CBB and agarose gel, respectively ^65^. The mononucleosome fractions were pooled, dialyzed against histone storage buffer, and H2AK119ub nucleosomes were purified by GST-UAB as described above, and cleaved from agarose beads with Thrombin (Millipore Sigma, T4648). The UAB-nucleosomes were further purified through a Superdex 200 gel filtration column (Cytiva, 28990994). Column fractions were analyzed with CBB and fractions containing mononucleosome fractions were concentrated to 3 mg/ml and subjected to Cryo-EM observation.

### Genome editing

For gene knockout, guidance RNAs (gRNAs) targeting human or mouse H2BC1, H2AZ.2, Ring1 (Ring1A), and RNF2 (Ring1B) were designed using the online software (https://chopchop.cbu.uib.no/), and the sequences were included in the respective figure panels. gRNAs and Cas9 plasmids were co-transfected into 293T or mouse R1 ESC cells with Lipofectamine^TM^ (Invitrogen, L3000015) or PolyJet^TM^ (SL100688, SignaGen Laboratories), respectively. Selection was made with puromycin for 3 days. Individual clones were picked up and expanded. The individual clones were first identified by genomic PCR, as shown the respective figures.

For gene knockin, multiple gRNAs targeting genomic regions flanking H2BC1 and H2AZ.2 transcription start sites were designed using the online software (https://chopchop.cbu.uib.no/), and the sequences were included in the respective figure panels. The efficiency for each gRNA in generating double-strand break was first evaluated with mCherry and GFP report system ^66^. The sgRNA with the highest efficiency was used for subsequent gene knockin experiments. Donor templates were constructed with 1000 bp of homologous arms on each side of the knockin site. The gRNA, donor template, and Cas9 were co-transfected into R1 cells with Polyjet. Selection was conducted with puromycin for 72 hrs, and the puromycin was removed after to reduce Cas9 genomic insertion. After individual clones were formed, these clones were hand-picked up and expanded to around 100 cells. Genomic DNA was purified with Tiangen Microgenomic DNA extraction Kit (Tiangen, DP316), and genomic PCR analyses were performed as described in each figure. The PCR products of the target regions were excised from gels and subjected to Sanger sequencing. Corrected targeted clones were further confirmed by immunoblots with antibody against the epitope tags.

### CUT&Tag and RNA-sequencing assay

The CUT&Tag assay was performed as previously described ^67^ using the Hyperactive Universal CUT&Tag Assay Kit for Illumina (Vazyme Biotech, #TD903). Briefly, 1 × 10^5^ cells were washed with 500 μl wash buffer and centrifuged at 2500 rpm for 5 min at room temperature. Cell pellets were resuspended with 100 μl wash buffer. A total of 10 μl concanavalin A-coated magnetic beads were washed twice with 100 μl binding buffer and then added to the cell tubes and incubated at room temperature for 15 minutes. After removing the supernatant, the bead-bound cells were resuspended with 50 μl antibody buffer (H2AK119ub and HA or flag diluted by 1:100). After incubation for overnight at 4°C, the primary antibody was carefully discarded, and pellets were then washed three times with Dig-wash buffer. Anti-rabbit IgG (vazyme, Ab207) diluted with 50 μl Dig-wash buffer was added to the cells. The cells were then incubated with rotation at room temperature for 1 h. The cells were then washed gently with 200 μl Dig-wash buffer three times, and 2 μl pA/G–Tnp together with 98 μl Dig-300 buffer was added to the samples. After incubating at room temperature for 1 h, the samples were washed gently with 200 μl Dig-300 buffer three times. Then, 10 μl 5 × TTBL mixed with 40 μl Dig-300 buffer was added to each sample, and the samples were incubated at 37 °C for 1 hr. The interactions were quenched by adding 5 μl 20 mg/ml Proteinase K, 100 μl Buffer L/B, and 20 μl DNA extraction beads and incubating the samples at 55°C for 10 min. The supernatant was discarded, and the beads were washed once with 200 μl Buffer WA and twice with 200 μl Buffer WB and resuspended with 22 μl nuclease free water. For library amplification, 15 μl of purified DNA was mixed with 25 μl of 2× CAM, along with 5 μl of uniquely barcoded i5 and i7 primers from the TruePrep Index Kit V2 for Illumina (Vazyme Biotech, # TD202). A total volume of 50 μl of sample was placed in a thermal cycler using the following program: 72 °C for 3 min; 98 °C for 3 min; 12–16 cycles of 98 °C for 10 s, 60 °C for 5 s, and 72°C for 1 min; and holding at 12 °C. To purify the PCR products, 2× volumes of VAHTS DNA Clean Beads (Vazyme Biotech, #N411) were added and incubated at room temperature for 5 minutes. The beads were washed twice with 200 μl fresh 80% ethanol and eluted in 22 μl ddH2O. All CUT&Tag libraries were sequenced by Novogene using the Illumina NovaSeq 6000 platform in PE150 mode (Novogene, Beijing, China).

### CUT&Tag and RNA-sequencing data analyses

Sequencing data files were first assessed for quality using FastQC v 0.11.9 (http://www.bioinformatics.babraham.ac.uk/projects/fastqc), and processed to remove residual adapter sequences with Cutadapt ^68^. Low-quality reads and bases were removed using Prinseq v 0.20.4. Resulting reads were aligned to the UCSC annotation of the mm10 mouse genome assembly using STAR aligner (v 2.7.8a --outSAMattributes NH HI NM MD AS nM XS -- outFilterMultimapScoreRange 2 --winAnchorMultimapNmax 1000 –outFilterMultimapNmax 10000 --outFilterMismatchNmax 10 --outFilterScoreMinOverLread 0.5 -- outFilterMatchNminOverLread 0.5 --outReadsUnmapped Fastx) ^69^. Picard (http://broadinstitute.github.io/picard) was used to mark PCR duplicates, which were filtered prior to further analysis with Samtools ^70^. Reads were counted with the Subread FeaturesCounts package v 2.0.2 (http://subread.sourceforge.net) ^71^ and normalized with Deseq2 v 2.11.40.6 ^72^.

Differential gene expression analysis was performed in Deseq2 v 2.11.40.6. Genes containing < 10 reads across replicates were removed from analysis. Genes with log2 fold change of > 1 were considered as differentially expressed. Deeptools BamCoverage was used to generate stranded bigwig files, normalized with the Deseq2 derived scale factors ^73^.

To assess the potential roles in intergenic histone marks and variants in splicing, we calculated splicing index values for all expressed genes as defined ^74^ :

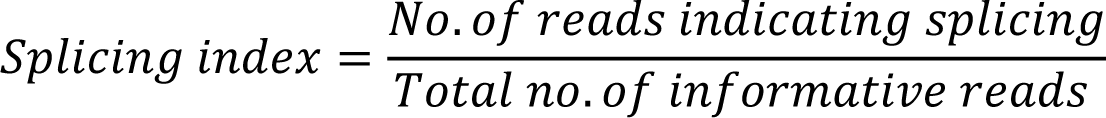

Briefly, aligned reads containing a cigar string with an “N” were considered indicative of a spliced read and were identified and counted with the FeatureCounts package. Next, reads containing both intron and exon derived sequences were identified and counted. Splicing index was calculated on a gene-by-gene basis using the above formula.

For Cut&Tag data, read quality was assessed with FastQC, and reads were preprocessed to remove adapter contamination, low-quality reads and low-quality bases as above. Reads passing quality control were mapped with to mm10 assembly of the mouse genome with the STAR aligner (--outFilterMultimapScoreRange 2 --winAnchorMultimapNmax 1000 --alignIntronMax 1 --alignMatesGapMax 1000 --outFilterMultimapNmax 10000 --outFilterMismatchNmax 10 -- outReadsUnmapped Fastx --alignEndsType Local --outSAMprimaryFlag AllBestScore). Duplicate reads were marked and excluded from subsequent analysis. Enriched regions were identified using the SEACR program ^75^, following the recommended workflow (https://github.com/FredHutch/SEACR). In brief, fragment size filtered reads were used to create bedgraph files, which were then used as input for the SEACR peak calling algorithm (0.01 norm stringent). Bedtools was used to aggregate peak lists across replicates to create a merged peak list used in subsequent analyses. Merged peaks were annotated in Homer ^76^. Heatmaps were generated with Deeptools on normalized input files ^73^.

For MNase-seq alignment and data analysis, reads were assessed for quality and preprocesses as above. Resulting reads were aligned to the UCSC annotation of the mm10 assembly of the mouse genome with STAR (--outFilterMultimapScoreRange 2 --winAnchorMultimapNmax 1000 --alignIntronMax 1 --alignMatesGapMax 1000 --outFilterMultimapNmax 10000 -- outFilterMismatchNmax 10). Duplicate reads were marked with Picard and filtered with Samtools. Multiply mapped reads and reads outside of the standard nucleosome size distribution (140bp – 160bp) were also removed prior to subsequent analysis. Peak calling was performed with Danpos ^77^ dtriple (--height 120). Peak annotation and motif identification was performed with Homer ^76^.

### Western blot

Whole cell extracts were obtained with dissolving cells in denature buffer (20 mM Tris, pH 7.4, 50 mM NaCl, 0.5% Nonidet P-40, 0.5% deoxycholate, 0.5% SDS, 1 mM EDTA) supplemented with protease inhibitor cocktail (Roche, Cat#4693116001). The lysates concentration was quantified by the BCA protein assay kit (Thermo Fisher Science, Cat#A53225). Cell lysates (30 μg) were loaded onto a 10% SDS-PAGE and transferred to a nitrocellulose membrane (Millipore, Cat#Z746010). The membrane was blocked with 5% (w/v) nonfat dry milk in TBST (0.1% Tween-20) and then incubated with the primary antibodies at 4°C for over-night, HRP-conjugated second anti-bodies at room temperature for 1 hr, and developed by Immobilon Western Chemiluminescent HRP Substrate (Millipore, Cat#WBKLS0500). Antibodies used are anti-HA (1:10000, Proteintech, Cat#51064-2-AP), anti-H3 (1:5000, Abcam, Cat#1791), anti-Flag (1:10000, Proteintech, Cat#20543-1-AP), anti-α-Tubulin (1:5000, Proteintech, Cat#11224-1-AP), anti-GAPDH (Fisher, PA1987), and anti-H2AK119ub (Cell signaling Technology 5019).

### Mass spectrometry

The intimal identification of H2BC1 and H2AZ.2 in H2AK119ub nucleosomes was carried out at UAB mass spectrometry facility. Briefly, Input and GST-pulldown nucleosomes were separated by SDS Bis-Tris gel (4–12%, Invitrogen). The gels were stained with SyproRuby. The entire lane for each sample was partitioned into 6MW fractions, and each gel plug was equilibrated in 100mM ammonium bicarbonate (AmBc). Each gel plug was then digested with Trypsin Gold (Promega) following the manufacturer’s instruction, and peptide extracts were reconstituted in 0.1% formic acid/ dd H2O at ∼0.1 μg/μL. Peptide digests were injected onto a 1260 Infinity nHPLC stack (Agilent Technologies) and separated using a 75-micron I.D. × 15 cm pulled tip C-18 column (Jupiter C-18 300 Å, 5 micron, Phenomenex). This system ran in-line with a Thermo Orbitrap Velos Pro hybrid mass spectrometer equipped with a nano-electrospray source (Thermo Fisher Scientific), and all data were collected in CID mode. The nHPLC was configured with binary mobile phases that included solvent A (0.1%FA in ddH2O), and solvent B (0.1%FA in 15% ddH2O / 85% ACN), programmed as follows: 10 min @ 5%B (2 μL/min, load), 90 min @ 5%–40%B (linear: 0.5 nL/min, analyze), 5 min @ 70%B (2 μL/min, wash), 10 min @ 5%B (2 μL/min, equilibrate). Following each parent ion scan (300-1200 m/z @ 60k resolution), fragmentation data (MS2) were collected on the top most intense 15 ions. For data dependent scans, charge state screening and dynamic exclusion were enabled with a repeat count of 2, repeat duration of 30 s, and exclusion duration of 90 s. The data were analyzed by database searching using the Sequest HT search algorithm with semi-tryptic digest using a custom All histone database downloaded from Swiss Prot. The data was processed with Scaffold v.5.3.2 with Scaffold PTM v.4.0.2 (Proteome Software). Searches included PTM’s of Oxidation of M, Acetylation of KR, methylation of KR, dimethylation of KR and GG mod on K (ubiquitination tag with tryptic digest).

The confirmation of H2BC1 and H2AZ.2 in H2AK119ub nucleosomes was carried out in the VCU Massey Comprehensive Cancer Center Proteomics Core. Briefly, protein bands containing histones were excised from the gel and cut into equal size cubes (approximately 1mm) then transferred to a siliconized Eppendorf tube. The gel pieces were washed with 80% acetonitrile ACN for 10 minutes, followed by an H2O wash for 10 minutes. Gel pieces were then dehydrated with 50% ACN for 5 minutes, then vacuum centrifuged for 20 minutes. 100ul of 1:50 dilution of acetylation reagent was added to the dried gel pieces and incubated for 1 hr. at 30°C with shaking at 1000 rpm. This step was repeated one more time from water wash to acetylation reaction to ensure higher acetylation efficiency. The supernatant was then discarded and gel pieces were washed twice with 200μl H_2_0 for 10 min with shaking. The samples were then dehydrated with 50% ACN for 5 minutes, then vacuum centrifuged for 20 min. The gel cubes were then incubated for 10 minutes in 40 μl trypsin (20μl of 12.5 ng/μl trypsin + 20 μl 100mM ammonium bicarbonate). The gel pieces were then overlaid with 20μl of 100mM ammonium bicarbonate and allowed to digest for 2 hrs at 37℃. The digests were collected into fresh siliconized tubes and centrifuged at 10,000 rpm for 5 minutes and then transferred to a fresh tube and stored for subsequent MS analysis. MS analyses were carried out using a Q Exactive HFX (Thermo) coupled to an Easy nLC 1200 (Thermo). The LC-MS system consisted of a Thermo Electron Q-Exactive HF-X mass spectrometer with an Easyspray Ion source connected to an Acclaim PepMap 75μm x 2cm nanoviper C18 3μm x 100Å pre-column in series with an Acclaim PepMap RSLC 75μm x 50cm C18 2μm bead size (Thermo Scientific). Peptides were injected into the column above. Peptides were eluted from the column with an 80%ACN to 0.1% formic acid gradient at a flow rate of 0.3μl/min over 1 hr. The nano-spray ion source was operated at 1.9 kV. The digests were analyzed using the rapid switching capability of the instrument thus acquiring a full scan mass spectrum to determine peptide molecular weights followed by product spectra (25 High Energy C-trap Dissociation HCD spectra). This mode of analysis produces approximately 50,000 MS/MS spectra of ions ranging in abundance over several orders of magnitude. Not all MS/MS spectra are derived from peptides.

### Cryo-EM data collection, image processing, and model building

The preparation of cryo-EM grids for nucleosomes involved the utilization of Vitrobot Mark IV (Thermo Fisher Scientific). A 4µl portion of the sample was applied onto individual Quantifoil holy carbon grids (R2/1, 300 mesh gold). Following a 4-second incubation at 8°C under 100% humidity, each grid underwent blotting and was subsequently plunged into liquid ethane at - 170°C. The data collection took place on a 300 kV Titan Krios microscope (Thermo Fisher Scientific) furnished with a Bioquantum Energy Filter and a K3 Summit direct electron detector (Gatan) at the Yale CCMI Electron Microscopy Facility. A comprehensive set of 5,630 movies was recorded, and further specifics can be found in Supplementary Table 1. For our datasets, we employed cryoSPARC v4 ^78^ for all subsequent data processing steps. Initially, about 4,887,804 particles were extracted from the images, followed by multiple rounds of 2D classification to isolate the highest-quality particles. Using the crystal structure 1AOI as the initial model, we proceeded with subsequent high-resolution 3D refinement. Several iterations of heterogeneous refinement were conducted on the top candidate models. The model with the highest resolution was selected for further refinement, aiming for a near-atomic resolution map. To progressively enhance the maps, a range of refinement techniques was employed. These included homogeneous refinement, local alignment refinement utilizing multiple masks for distinct regions, optimization of high-order attenuation parameters for individual optical groups, and local CTF refinement, similar to our previous description^79^. Post-refinement, particle coordinates were re-centered and re-extracted from the original micrographs, producing a new particle set comprising the final count of 1,288,371 particles. As a result, the majority of mastigoneme filament regions were refined to approximately 2.6 Å resolution, facilitating the construction of an atomic model with assigned side chains in Coot ^80^ (See Fig. 2). The majority of regions were refined to a resolution finer than 2.6 Å, enabling the construction of atomic models with assigned side chains using Coot. The final model underwent automatic refinement using Phenix and Isolde, with iterations repeated until all parameters reached a satisfactory level of refinement. Visual representations in figures and movies were generated using ChimeraX ^81^.

### Experiments in *Xenopus laevis* embryos

*Xenopus laevis* embryos were obtained using standard procedures ^82,83^ approved by the VCU Institutional Animal Care and Use Committee (IACUC protocol number AD20261). Embryos were staged according to Nieuwkoop and Faber ^84,85^. Stages are also reported as hours post fertilization at 23°C. To knockdown gene function we used antisense oligonucleotides stabilized with morpholino rings (Morpholinos (MOs) GeneTools). Sequences and details available upon request. A standard control MO that does not target any Xenopus sequence was utilized as a negative control. Embryos were injected with a Femtojet injection system using standard practices ^86^. After injections, embryos were placed into 10 cm culture dishes filled with frog embryo media and maintained in a 15°C incubator. Media was changed daily until embryos reached a swimming larval stage when they were fixed in 4% paraformaldehyde and imaged on a Zeiss stereoscope fitted with an AxioCam digital camera.

## Acknowledgments

The authors would like to express their gratitude to Dr. Terje Dokland (University of Alabama at Birmingham) for preliminary cryo-EM studies, Dr. Chenbei Chang and Mr. Saeid Mohammad Parast (University of Alabama at Birmingham) for preliminary *Xenopus* studies, Dr. Robert McGinty (University of North Carolina at Chapel Hills) for reagents and consultation of structural studies, and Drs. Brian D Strahl (University of North Carolina at Chapel Hills), Louise Chow (University of Alabama at Birmingham), Dr. Mitzi Kuroda (Harvard University) for suggestions and discussions. H.W. is supported by a start-up package from VCU, grants from NIH (GM130696), and DOD (W81XWH-21-1-0714). K.Z. is supported by an NIH grant (R35GM142959). VCU Massey Comprehensive Cancer Center Proteomics Shared Resource is supported, in part, with funding from NIH-NCI Cancer Center Support Grant P30 CA016059. Jiahuan Zheng is supported by Bau Tsu Zung Bau Kwan Yeu Hing Research and Clinical Fellowship from the University of Hong Kong.

## Authors contributions

H.W. conceived the concept and guided all the studies. X.S., C.C., Y.W., W. Z., J.Z, and H.Z. performed experiments. A.J and B.Z. analyzed the CUT&RUN and RNA-sequencing data. J.M., C.L., and A.R. performed the mass spectrometry study. K.Z. supervised the cryo-EM data collection, image processing, and model building. G.C. supervised the knockin and knockout studies in mouse ESC. A.D. performed experiments with *Xenopus*. H.W., K.Z, A.D., and R.W. wrote the manuscript with contributions from all authors.

## Data Availability

All high-throughput sequencing data have been deposited to NCBI GEO repository with accession number GSE xxxx. The Cryo-EM data has been deposited in the Protein Data Bank, www.PDB.org (PBD ID 8VLR, EMD43342).

## Bibliography

1 Wang, H. et al. Role of histone H2A ubiquitination in Polycomb silencing. Nature 431, 873–878, doi:10.1038/nature02985 (2004).

2 Barbour, H., Daou, S., Hendzel, M. & Affar, E. B. Polycomb group-mediated histone H2A monoubiquitination in epigenome regulation and nuclear processes. Nature communications 11, 5947, doi:10.1038/s41467-020-19722-9 (2020).

3 Vassilev, A. P., Rasmussen, H. H., Christensen, E. I., Nielsen, S. & Celis, J. E. The levels of ubiquitinated histone H2A are highly upregulated in transformed human cells: partial colocalization of uH2A clusters and PCNA/cyclin foci in a fraction of cells in S-phase. Journal of cell science 108 (Pt 3), 1205–1215, doi:10.1242/jcs.108.3.1205 (1995).

4 Lee, H. S., Lee, S. A., Hur, S. K., Seo, J. W. & Kwon, J. Stabilization and targeting of INO80 to replication forks by BAP1 during normal DNA synthesis. Nature communications 5, 5128, doi:10.1038/ncomms6128 (2014).

5 Lee, H. S. et al. BAP1 promotes stalled fork restart and cell survival via INO80 in response to replication stress. The Biochemical journal 476, 3053–3066, doi:10.1042/bcj20190622 (2019).

6 Bergink, S. et al. DNA damage triggers nucleotide excision repair-dependent monoubiquitylation of histone H2A. Genes & development 20, 1343–1352, doi:10.1101/gad.373706 (2006).

7 Chou, D. M. et al. A chromatin localization screen reveals poly (ADP ribose)-regulated recruitment of the repressive polycomb and NuRD complexes to sites of DNA damage. Proceedings of the National Academy of Sciences of the United States of America 107, 18475–18480, doi:10.1073/pnas.1012946107 (2010).

8 Ismail, I. H., Andrin, C., McDonald, D. & Hendzel, M. J. BMI1-mediated histone ubiquitylation promotes DNA double-strand break repair. The Journal of cell biology 191, 45–60, doi:10.1083/jcb.201003034 (2010).

9 Ginjala, V. et al. BMI1 is recruited to DNA breaks and contributes to DNA damage-induced H2A ubiquitination and repair. Molecular and cellular biology 31, 1972–1982, doi:10.1128/mcb.00981-10 (2011).

10 Ismail, I. H. et al. CBX4-mediated SUMO modification regulates BMI1 recruitment at sites of DNA damage. Nucleic acids research 40, 5497–5510, doi:10.1093/nar/gks222 (2012).

11 de Napoles, M. et al. Polycomb group proteins Ring1A/B link ubiquitylation of histone H2A to heritable gene silencing and X inactivation. Developmental cell 7, 663–676, doi:10.1016/j.devcel.2004.10.005 (2004).

12 Fang, J., Chen, T., Chadwick, B., Li, E. & Zhang, Y. Ring1b-mediated H2A ubiquitination associates with inactive X chromosomes and is involved in initiation of X inactivation. The Journal of biological chemistry 279, 52812–52815, doi:10.1074/jbc.C400493200 (2004).

13 Bernstein, E. et al. Mouse polycomb proteins bind differentially to methylated histone H3 and RNA and are enriched in facultative heterochromatin. Molecular and cellular biology 26, 2560–2569, doi:10.1128/mcb.26.7.2560-2569.2006 (2006).

14 Bravo, M. et al. Polycomb RING1A- and RING1B-dependent histone H2A monoubiquitylation at pericentromeric regions promotes S-phase progression. Journal of cell science 128, 3660–3671, doi:10.1242/jcs.173021 (2015).

15 Goldknopf, I. L. et al. Isolation and characterization of protein A24, a “histone-like” non-histone chromosomal protein. The Journal of biological chemistry 250, 7182–7187 (1975).

16 Piunti, A. & Shilatifard, A. Epigenetic balance of gene expression by Polycomb and COMPASS families. *Science (New York*, N.Y.) 352, aad9780, doi:10.1126/science.aad9780 (2016).

17 Schuettengruber, B., Bourbon, H. M., Di Croce, L. & Cavalli, G. Genome Regulation by Polycomb and Trithorax: 70 Years and Counting. Cell 171, 34–57, doi:10.1016/j.cell.2017.08.002 (2017).

18 Blackledge, N. P. & Klose, R. J. The molecular principles of gene regulation by Polycomb repressive complexes. Nature reviews. Molecular cell biology 22, 815–833, doi:10.1038/s41580-021-00398-y (2021).

19 Piunti, A. & Shilatifard, A. The roles of Polycomb repressive complexes in mammalian development and cancer. Nature Reviews Molecular Cell Biology 22, 326–345, doi:10.1038/s41580-021-00341-1 (2021).

20 Doyle, E. J., Morey, L. & Conway, E. Know when to fold ’em: Polycomb complexes in oncogenic 3D genome regulation. Front Cell Dev Biol 10, 986319, doi:10.3389/fcell.2022.986319 (2022).

21 Pengelly, A. R., Kalb, R., Finkl, K. & Müller, J. Transcriptional repression by PRC1 in the absence of H2A monoubiquitylation. Genes & development 29, 1487–1492, doi:10.1101/gad.265439.115 (2015).

22 Francis, N. J., Kingston, R. E. & Woodcock, C. L. Chromatin compaction by a polycomb group protein complex. *Science (New York*, N.Y.) 306, 1574–1577, doi:10.1126/science.1100576 (2004).

23 Eskeland, R. et al. Ring1B compacts chromatin structure and represses gene expression independent of histone ubiquitination. Molecular cell 38, 452–464, doi:10.1016/j.molcel.2010.02.032 (2010).

24 Grau, D. J. et al. Compaction of chromatin by diverse Polycomb group proteins requires localized regions of high charge. Genes & development 25, 2210–2221, doi:10.1101/gad.17288211 (2011).

25 Schoenfelder, S. et al. Polycomb repressive complex PRC1 spatially constrains the mouse embryonic stem cell genome. Nature genetics 47, 1179–1186, doi:10.1038/ng.3393 (2015).

26 Lau Mei, S., et al. Mutation of a nucleosome compaction region disrupts Polycomb-mediated axial patterning. *Science (New York*, N.Y.) 355, 1081–1084, doi:10.1126/science.aah5403 (2017).

27 Tsuboi, M. et al. Ubiquitination-Independent Repression of PRC1 Targets during Neuronal Fate Restriction in the Developing Mouse Neocortex. Developmental cell 47, 758–772.e755, doi:10.1016/j.devcel.2018.11.018 (2018).

28 Endoh, M. et al. Histone H2A mono-ubiquitination is a crucial step to mediate PRC1-dependent repression of developmental genes to maintain ES cell identity. PLoS genetics 8, e1002774, doi:10.1371/journal.pgen.1002774 (2012).

29 Blackledge, N. P. et al. Variant PRC1 complex-dependent H2A ubiquitylation drives PRC2 recruitment and polycomb domain formation. Cell 157, 1445–1459, doi:10.1016/j.cell.2014.05.004 (2014).

30 Fursova, N. A. et al. Synergy between Variant PRC1 Complexes Defines Polycomb-Mediated Gene Repression. Molecular cell 74, 1020–1036.e1028, doi:10.1016/j.molcel.2019.03.024 (2019).

31 Blackledge, N. P. et al. PRC1 Catalytic Activity Is Central to Polycomb System Function. Molecular cell 77, 857–874.e859, doi:10.1016/j.molcel.2019.12.001 (2020).

32 Tamburri, S. et al. Histone H2AK119 Mono-Ubiquitination Is Essential for Polycomb-Mediated Transcriptional Repression. Molecular cell 77, 840–856.e845, doi:10.1016/j.molcel.2019.11.021 (2020).

33 Dobrinić, P., Szczurek, A. T. & Klose, R. J. PRC1 drives Polycomb-mediated gene repression by controlling transcription initiation and burst frequency. Nature structural & molecular biology 28, 811–824, doi:10.1038/s41594-021-00661-y (2021).

34 Lee, H. G., Kahn, T. G., Simcox, A., Schwartz, Y. B. & Pirrotta, V. Genome-wide activities of Polycomb complexes control pervasive transcription. Genome research 25, 1170–1181, doi:10.1101/gr.188920.114 (2015).

35 Scelfo, A. et al. Functional Landscape of PCGF Proteins Reveals Both RING1A/B-Dependent-and RING1A/B-Independent-Specific Activities. Molecular cell 74, 1037–1052.e1037, doi:10.1016/j.molcel.2019.04.002 (2019).

36 Conway, E. et al. BAP1 enhances Polycomb repression by counteracting widespread H2AK119ub1 deposition and chromatin condensation. Molecular cell 81, 3526–3541.e3528, doi:10.1016/j.molcel.2021.06.020 (2021).

37 Fursova, N. A. et al. BAP1 constrains pervasive H2AK119ub1 to control the transcriptional potential of the genome. Genes & development 35, 749–770, doi:10.1101/gad.347005.120 (2021).

38 Scheuermann, J. C. et al. Histone H2A deubiquitinase activity of the Polycomb repressive complex PR-DUB. Nature 465, 243–247, doi:10.1038/nature08966 (2010).

39 Balasubramani, A. et al. Cancer-associated ASXL1 mutations may act as gain-of-function mutations of the ASXL1-BAP1 complex. Nature communications 6, 7307, doi:10.1038/ncomms8307 (2015).

40 Gao, Z. et al. An AUTS2-Polycomb complex activates gene expression in the CNS. Nature 516, 349–354, doi:10.1038/nature13921 (2014).

41 Pivetti, S. et al. Loss of PRC1 activity in different stem cell compartments activates a common transcriptional program with cell type-dependent outcomes. Science advances 5, eaav1594, doi:10.1126/sciadv.aav1594 (2019).

42 Nakagawa, T. et al. Deubiquitylation of histone H2A activates transcriptional initiation via trans-histone cross-talk with H3K4 di- and trimethylation. Genes & development 22, 37–49, doi:10.1101/gad.1609708 (2008).

43 Zhou, W. et al. Histone H2A monoubiquitination represses transcription by inhibiting RNA polymerase II transcriptional elongation. Molecular cell 29, 69–80, doi:10.1016/j.molcel.2007.11.002 (2008).

44 Wang, Y. Z. et al. H2A mono-ubiquitination differentiates FACT’s functions in nucleosome assembly and disassembly. Nucleic acids research 50, 833–846, doi:10.1093/nar/gkab1271 (2022).

45 Jason, L. J., Finn, R. M., Lindsey, G. & Ausió, J. Histone H2A ubiquitination does not preclude histone H1 binding, but it facilitates its association with the nucleosome. The Journal of biological chemistry 280, 4975–4982, doi:10.1074/jbc.M410203200 (2005).

46 Zhu, P. et al. A histone H2A deubiquitinase complex coordinating histone acetylation and H1 dissociation in transcriptional regulation. Molecular cell 27, 609–621, doi:10.1016/j.molcel.2007.07.024 (2007).

47 Kalb, R. et al. Histone H2A monoubiquitination promotes histone H3 methylation in Polycomb repression. Nature structural & molecular biology 21, 569–571, doi:10.1038/nsmb.2833 (2014).

48 Cooper, S. et al. Jarid2 binds mono-ubiquitylated H2A lysine 119 to mediate crosstalk between Polycomb complexes PRC1 and PRC2. Nature communications 7, 13661, doi:10.1038/ncomms13661 (2016).

49 Cooper, S. et al. Targeting polycomb to pericentric heterochromatin in embryonic stem cells reveals a role for H2AK119u1 in PRC2 recruitment. Cell reports 7, 1456–1470, doi:10.1016/j.celrep.2014.04.012 (2014).

50 Cao, R. et al. Role of histone H3 lysine 27 methylation in Polycomb-group silencing. *Science (New York*, N.Y.) 298, 1039–1043, doi:10.1126/science.1076997 (2002).

51 Boyle, S. et al. A central role for canonical PRC1 in shaping the 3D nuclear landscape. Genes & development 34, 931–949, doi:10.1101/gad.336487.120 (2020).

52 Zhang, Z. et al. Role of remodeling and spacing factor 1 in histone H2A ubiquitination-mediated gene silencing. Proceedings of the National Academy of Sciences of the United States of America 114, E7949–e7958, doi:10.1073/pnas.1711158114 (2017).

53 Horikoshi, N. et al. Structural polymorphism in the L1 loop regions of human H2A.Z.1 and H2A.Z.2. *Acta crystallographica*. Section D, Biological crystallography 69, 2431–2439, doi:10.1107/s090744491302252x (2013).

54 Martire, S. & Banaszynski, L. A. The roles of histone variants in fine-tuning chromatin organization and function. Nature reviews. Molecular cell biology 21, 522–541, doi:10.1038/s41580-020-0262-8 (2020).

55 Ge, W. et al. Basis of the H2AK119 specificity of the Polycomb repressive deubiquitinase. Nature 616, 176–182, doi:10.1038/s41586-023-05841-y (2023).

56 Thomas, J. F. et al. Structural basis of histone H2A lysine 119 deubiquitination by Polycomb repressive deubiquitinase BAP1/ASXL1. Science Advances 9, eadg9832, doi:doi:10.1126/sciadv.adg9832 (2023).

57 Montellier, E. et al. Chromatin-to-nucleoprotamine transition is controlled by the histone H2B variant TH2B. Genes & development 27, 1680–1692, doi:10.1101/gad.220095.113 (2013).

58 Shinagawa, T. et al. Histone Variants Enriched in Oocytes Enhance Reprogramming to Induced Pluripotent Stem Cells. Cell stem cell 14, 217–227, 10.1016/j.stem.2013.12.015 (2014).

59 Huynh, L. M., Shinagawa, T. & Ishii, S. Two Histone Variants TH2A and TH2B Enhance Human Induced Pluripotent Stem Cell Generation. Stem Cells Dev 25, 251–258, doi:10.1089/scd.2015.0264 (2016).

60 Creyghton, M. P. et al. H2AZ is enriched at polycomb complex target genes in ES cells and is necessary for lineage commitment. Cell 135, 649–661, doi:10.1016/j.cell.2008.09.056 (2008).

61 Padavattan, S. et al. Structural and functional analyses of nucleosome complexes with mouse histone variants TH2a and TH2b, involved in reprogramming. Biochemical and biophysical research communications 464, 929–935, doi:10.1016/j.bbrc.2015.07.070 (2015).

62 Padavattan, S. et al. Structural analyses of the nucleosome complexes with human testis-specific histone variants, hTh2a and hTh2b. Biophysical chemistry 221, 41–48, doi:10.1016/j.bpc.2016.11.013 (2017).

63 Żylicz, J. J., et al. The Implication of Early Chromatin Changes in X Chromosome Inactivation. Cell 176, 182–197.e123, doi:10.1016/j.cell.2018.11.041 (2019).

64 Yang, W. et al. The histone H2A deubiquitinase Usp16 regulates embryonic stem cell gene expression and lineage commitment. Nature communications 5, 3818, doi:10.1038/ncomms4818 (2014).

65 Jones, A., Joo, H. Y., Robbins, W. & Wang, H. Purification of histone ubiquitin ligases from HeLa cells. *Methods (San Diego*, Calif.) 54, 315–325, doi:10.1016/j.ymeth.2011.03.003 (2011).

66 Kim, H. et al. Surrogate reporters for enrichment of cells with nuclease-induced mutations. Nature methods 8, 941–943, doi:10.1038/nmeth.1733 (2011).

67 Kaya-Okur, H. S. et al. CUT&Tag for efficient epigenomic profiling of small samples and single cells. Nature communications 10, 1930, doi:10.1038/s41467-019-09982-5 (2019).

68 Schmieder, R. & Edwards, R. Quality control and preprocessing of metagenomic datasets. Bioinformatics (Oxford, England) 27, 863–864, doi:10.1093/bioinformatics/btr026 (2011).

69 Dobin, A. et al. STAR: ultrafast universal RNA-seq aligner. *Bioinformatics (Oxford*, England) 29, 15–21, doi:10.1093/bioinformatics/bts635 (2013).

70 Li, H. et al. The Sequence Alignment/Map format and SAMtools. *Bioinformatics (Oxford*, England) 25, 2078–2079, doi:10.1093/bioinformatics/btp352 (2009).

71 Liao, Y., Smyth, G. K. & Shi, W. featureCounts: an efficient general purpose program for assigning sequence reads to genomic features. *Bioinformatics (Oxford*, England) 30, 923–930, doi:10.1093/bioinformatics/btt656 (2014).

72 Love, M. I., Huber, W. & Anders, S. Moderated estimation of fold change and dispersion for RNA-seq data with DESeq2. Genome biology 15, 550, doi:10.1186/s13059-014-0550-8 (2014).

73 Ramírez, F. et al. deepTools2: a next generation web server for deep-sequencing data analysis. Nucleic acids research 44, W160–165, doi:10.1093/nar/gkw257 (2016).

74 Tilgner, H. et al. Deep sequencing of subcellular RNA fractions shows splicing to be predominantly co-transcriptional in the human genome but inefficient for lncRNAs. Genome research 22, 1616–1625, doi:10.1101/gr.134445.111 (2012).

75 Meers, M. P., Tenenbaum, D. & Henikoff, S. Peak calling by Sparse Enrichment Analysis for CUT&RUN chromatin profiling. Epigenetics & chromatin 12, 42, doi:10.1186/s13072-019-0287-4 (2019).

76 Heinz, S. et al. Simple combinations of lineage-determining transcription factors prime cis-regulatory elements required for macrophage and B cell identities. Molecular cell 38, 576–589, doi:10.1016/j.molcel.2010.05.004 (2010).

77 Chen, K. et al. DANPOS: dynamic analysis of nucleosome position and occupancy by sequencing. Genome research 23, 341–351, doi:10.1101/gr.142067.112 (2013).

78 Punjani, A., Rubinstein, J. L., Fleet, D. J. & Brubaker, M. A. cryoSPARC: algorithms for rapid unsupervised cryo-EM structure determination. Nature methods 14, 290–296, doi:10.1038/nmeth.4169 (2017).

79 Ton, W. D. et al. Microtubule-binding-induced allostery triggers LIS1 dissociation from dynein prior to cargo transport. Nature Structural & Molecular Biology, 1–15 (2023).

80 Emsley, P. & Cowtan, K. Coot: model-building tools for molecular graphics. Acta crystallographica section D: biological crystallography 60, 2126–2132 (2004).

81 Goddard, T. D. et al. UCSF ChimeraX: Meeting modern challenges in visualization and analysis. Protein Science 27, 14–25 (2018).

82 Sive, H., Grainger, R. & Harland, R. Early Development of Xenopus laevis: a laboratory manual. (Cold Spring Harbor Laboratory Press, 2000).

83 Sive, H. L., Grainger, R. M. & Harland, R. M. Xenopus laevis In Vitro Fertilization and Natural Mating Methods. CSH protocols 2007, pdb.prot4737, doi:10.1101/pdb.prot4737 (2007).

84 Nieuwkoop, P. & Faber, J. Normal table of Xenopus laevis (Daudin): a systematic and chronological survey of the development from the fertilized egg till the end of metamorphosis. (North-Holland, 1967).

85 Nieuwkoop, P. D. & Faber, J. Normal table of Xenopus laevis (Daudin) : a systematical and chronological survey of the development from the fertilized egg till the end of metamorphosis. (Garland Pub., 1994).

86 Sive, H. L., Grainger, R. M. & Harland, R. M. Microinjection of Xenopus embryos. Cold Spring Harbor protocols 2010, pdb.ip81, doi:10.1101/pdb.ip81 (2010).

87 Fisher, M. et al. Xenbase: key features and resources of the Xenopus model organism knowledgebase. Genetics 224, doi:10.1093/genetics/iyad018 (2023).

