## Supplemental file for "Role of histone variants H2BC1 and H2AZ.2 in H2AK119ub nucleosome organization and Polycomb gene silencing"

Supplemental Figure S1-S10 with legends

Supplemental Table 1

Supplementary Figure 1

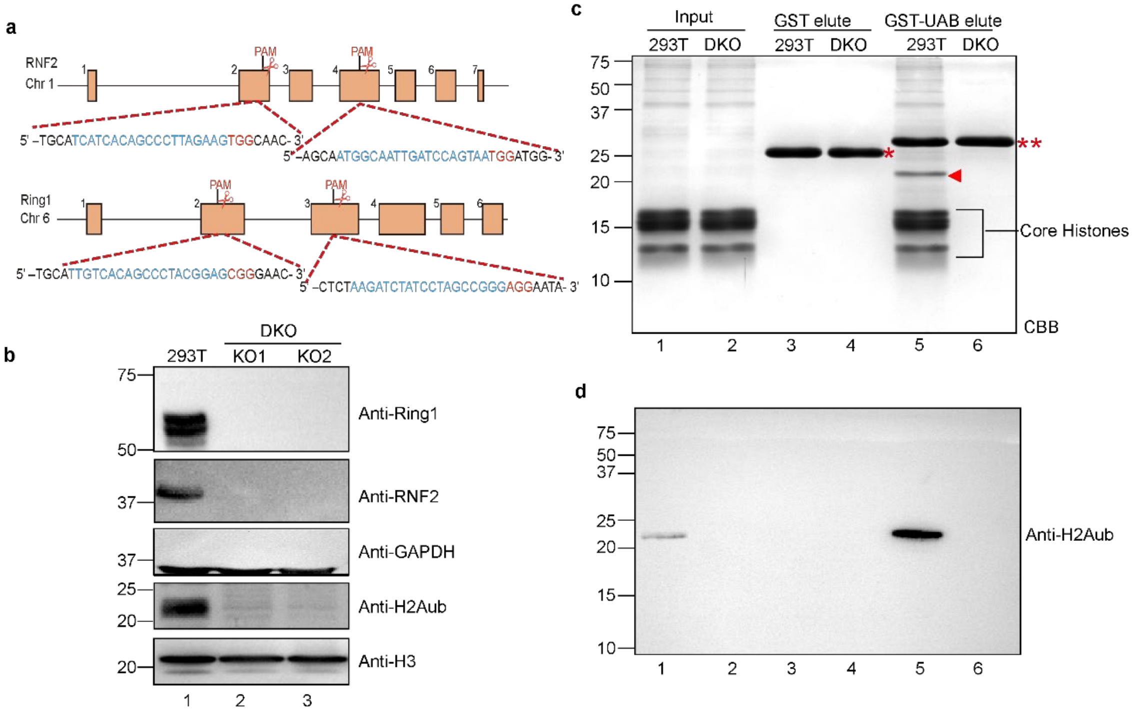


**Figure S1. GST-UAB pulled down H2AK119ub nucleosomes from control but not Ring1/Ring2 double knockout 293T cells.**

**a.** Schematic diagram of guide RNA (gRNA) design targeting human Ring1 and Ring2 (RNF2). The sgRNA-targeting sequences are in blue, and PAM sequences are in red.

**b.** Immunoblot assay of control and two Ring1 and RNF2 double knockout (DKO) clones. Antibodies used and molecular markers are labeled on the right and left side of the panel, respectively. GAPDH and histone H3 were used as loading controls.

**c-d**. Coomassie brilliant blue (CBB) staining (**c**) and anti-H2AK119ub immunoblots (**d**) of an SDS-PAGE containing an aliquot of Input (lanes 1, 2) and Bound nucleosomes in GST (lanes 3, 4) or GST-UAB (lanes 5, 6) pulldown assays. H2AK119ub is indicated with a red arrowhead. GST and GST-UAB are marked with a single and double red asterisk, respectively. Molecular markers are shown on the left. The positions of core histones are labeled.

### Supplementary Figure 2

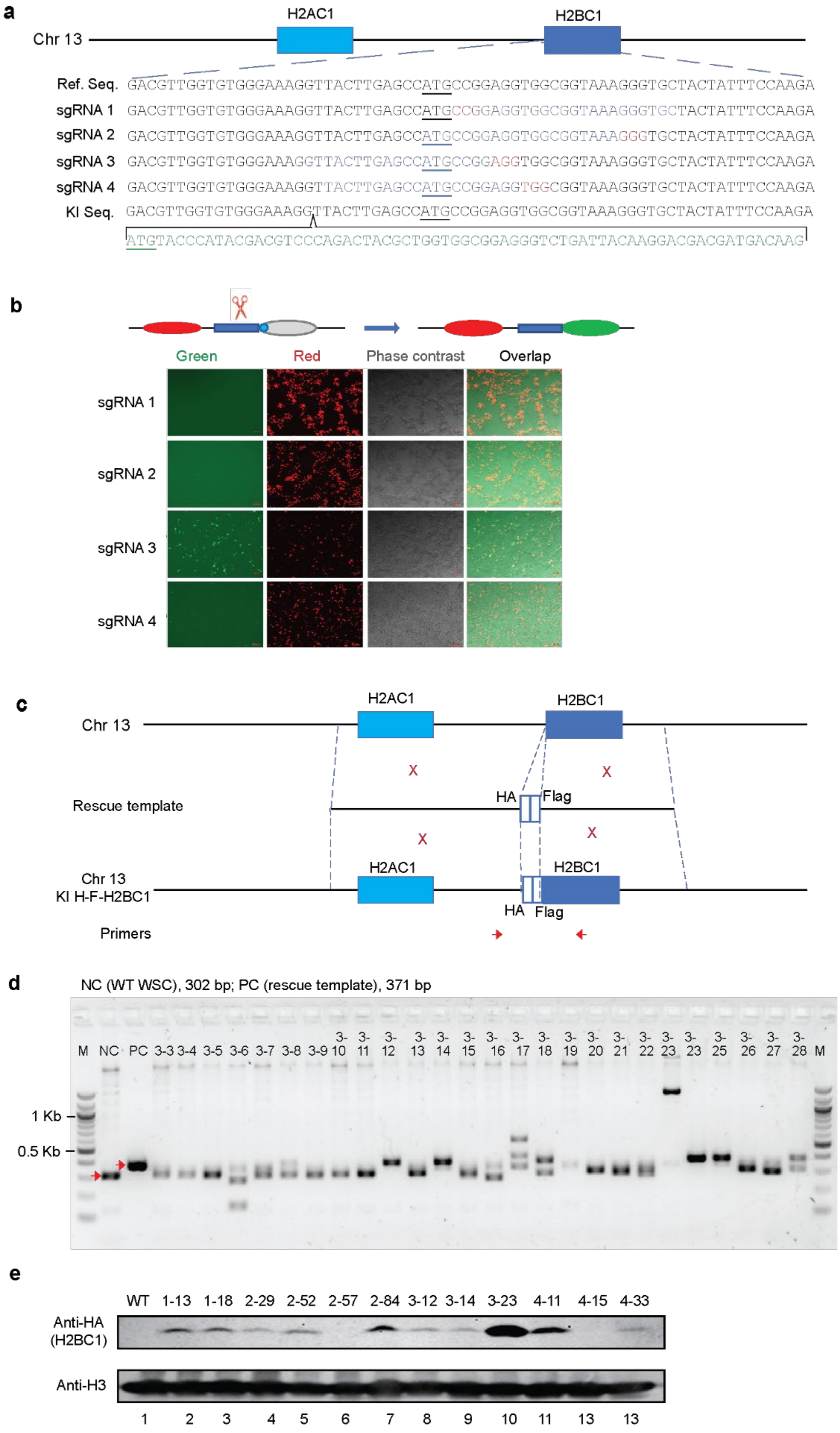

**Figure S2. The strategy for generating HA-Flag-H2BC1 knockin mouse ESC lines.**

- **a.** Schematic diagram of guide RNA (gRNA) design targeting H2BC1 gene regions flanking the translation start site ATG. The sgRNA-targeting sequences are in blue, PAM sequences are in red, and the translation start site ATG is underlined.
- **b.** GFP/mCherry assays to evaluate the efficiency of various H2BC1 gRNAs, as indicated panel **a,** in generating double-strand breaks.
- **c.** Schematic diagram of the strategy knocking a Flag-HA tag into the mouse H2BC1 gene. The rescue template contains 1000 bp homologous arms up- and down- stream of the translation start site ATG, respectively. Primers used to identify positive colonies are shown at the bottom. Sequences of the corrected targeted Flag-HA-H2BC1 KI ESC clones are shown in panel **a** (green).
- **d.** An agarose gel containing PCR products from wild-type (WT) mouse R1 ESC (NC, negative control), rescue template (PC, positive control), and individual single colonies from HA-Flag-H2BC1 KI experiments. Target bands are indicated with red arrows. A representative result is shown.
- **e.** Immunoblots of control (WT R1 ESC) and candidate HA-Flag-H2BC1 KI mouse ESC lines.

### Supplementary Figure 3

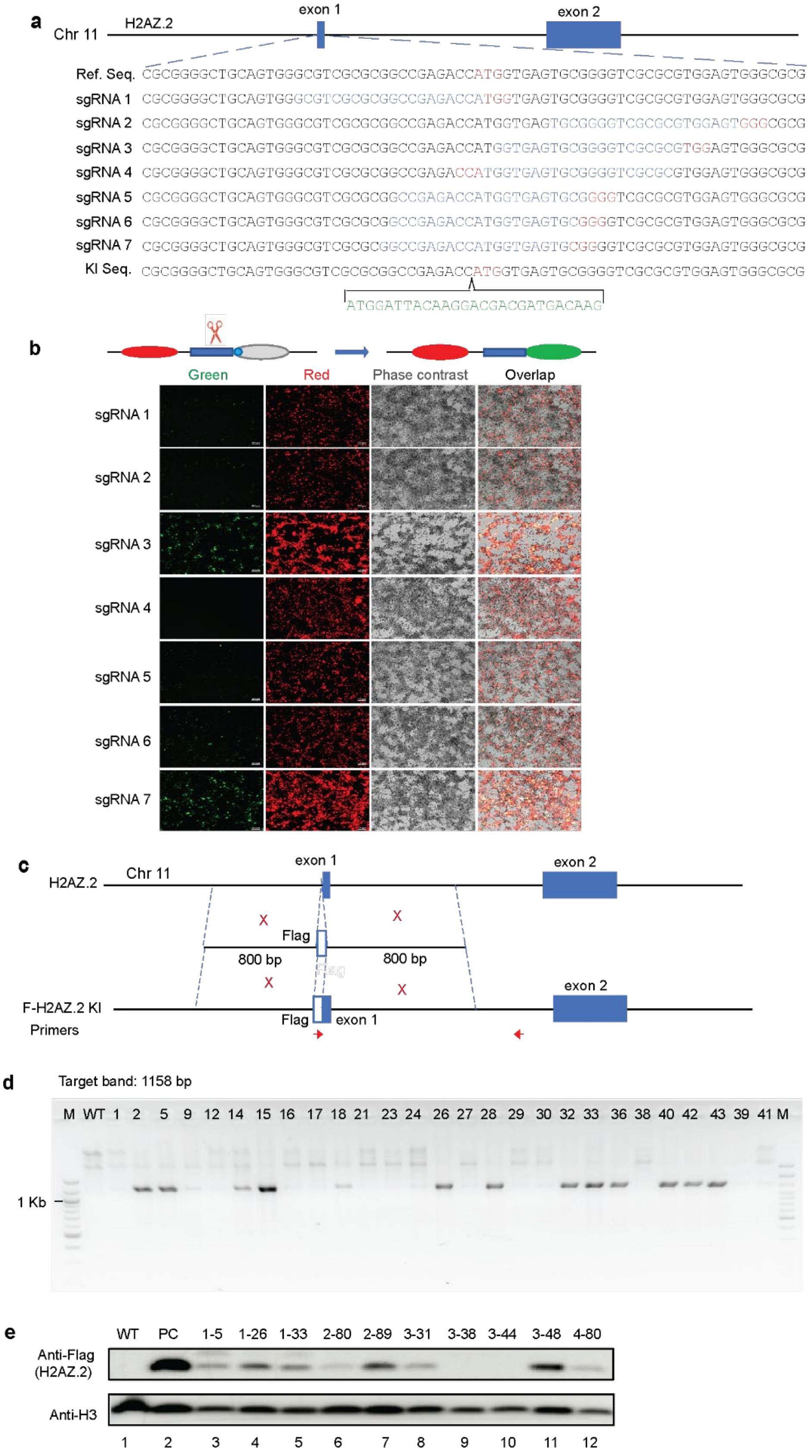

**Figure S3. The strategy for generating Flag-H2AZ.2 knockin mouse ESC lines.**

- **a.** Schematic diagram of guide RNA (gRNA) design targeting H2AZ.2 gene regions flanking the translation start site ATG. The sgRNA-targeting sequences are in blue, PAM sequences are in red, and the translation start site ATG is underlined.
- **b.** GFP/mCherry assays to evaluate the efficiency of various H2AZ.2 gRNAs, as indicated in panel **a,** in generating double-strand breaks.
- **c.** Schematic diagram of the strategy knocking a Flag tag into the mouse H2AZ.2 gene. The rescue template contains 1000 bp homologous arms up- and down- stream of the translation start site ATG, respectively. Primers used to identify positive colonies are shown at the bottom. Sequences of the corrected targeted Flag-H2AZ.2 KI ESC clones are shown in panel **a** (green).
- **d.** An agarose gel containing PCR products from wild-type (WT) mouse R1 ESC (NC, negative control), rescue template (PC, positive control), and individual single colonies from Flag-H2AZ.2 KI experiments. Target bands are indicated with red arrows. A representative result is shown.
- **e.** Immunoblots of control (WT R1 ESC) and candidate Flag-H2AZ.2 KI mouse ESC lines.

### Supplementary Figure 4

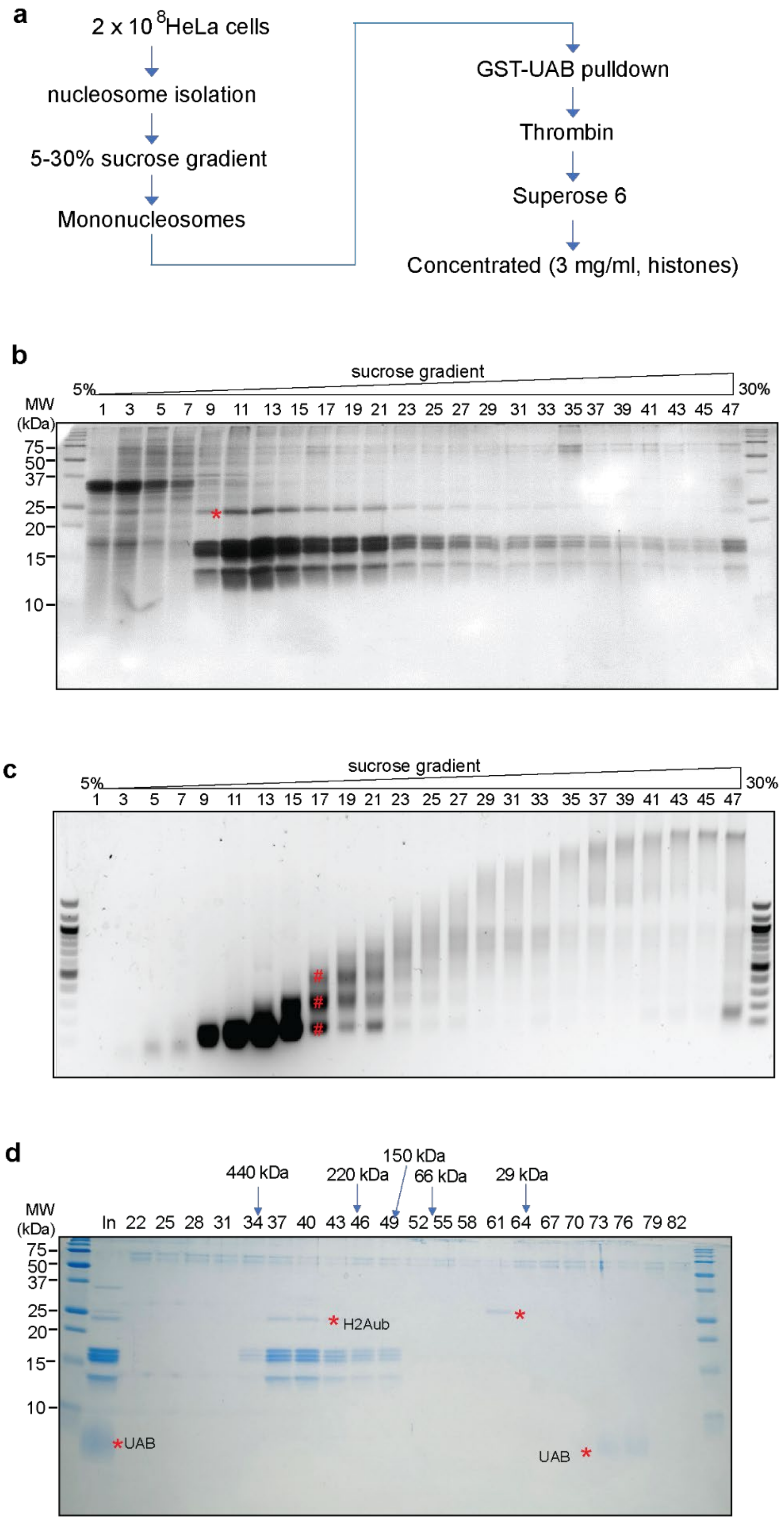

**Figure S4. Sample preparation of H2AK119ub nucleosomes for Cryo-EM.**

**a.** Schematic diagram of the experimental procedure.

**b-c.** CBB staining of an SDS-PAGE (**b**) and SYBR staining of an agarose gel (**c**) containing an aliquot of fractions of 5-30% sucrose gradient of nucleosomes purified from HeLa cells. H2AK119ub is denoted with *. Nucleosome positions are denoted with #.

**d.** CBB staining of an SDS-PAGE containing an aliquot of fractions of S200 gel filtration. GST-UAB bound nucleosomes were cleaved from agarose beads and subjected to gel filtration purification. The positions of H2AK119ub, GST, and UAB are labeled. The molecular weight is marked at the top of the fractions.

### Supplementary Figure 5

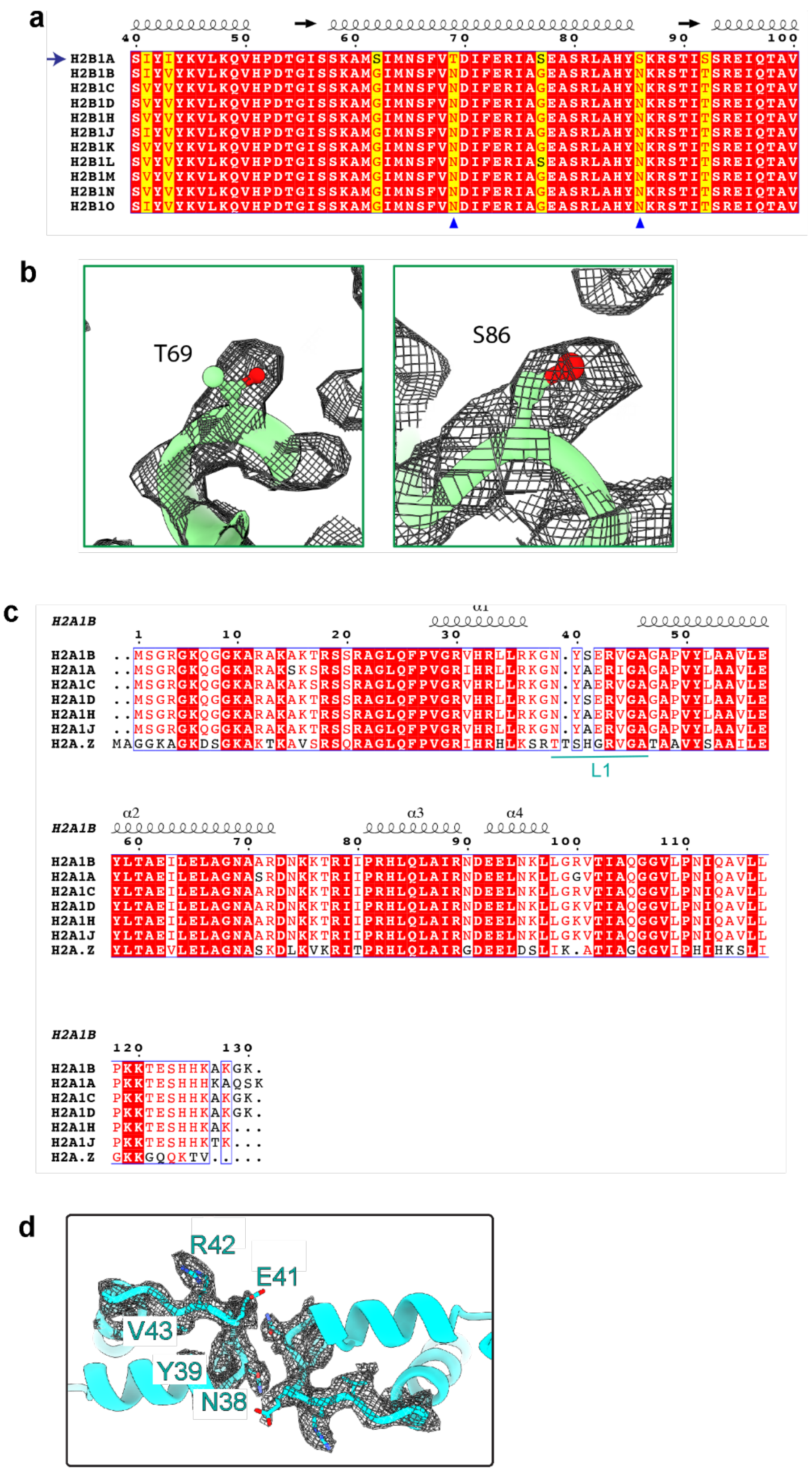

**Figure S5. Identification of H2BC1 and H2A1A in H2AK119ub nucleosomes.**

- **a.** Sequence alignment of H2BC1 (H2B1A) with other H2B isoform. Amino acids unique to H2BC1 were highlighted in orange. Two amino acids distinction H2BC1 from other isoforms are marked with arrowheads. The predicated second structure is shown in the top.
- **b.** Electronic density map of T69 and S86, which allows the identification of H2BC1 in H2AK119ub nucleosomes.
- **c.** Sequence alignment of H2A1B with the other H2A isoforms. Identical amino acids are in red. The predicated second structure is shown in the top. L1 loop, which allows the identification of H2A1.B, is marked.
- **d.** Electronic density map around the L1 loop of H2A, allowing the identification of H2A1B in H2AK119ub nucleosomes.

### Supplementary Figure 6

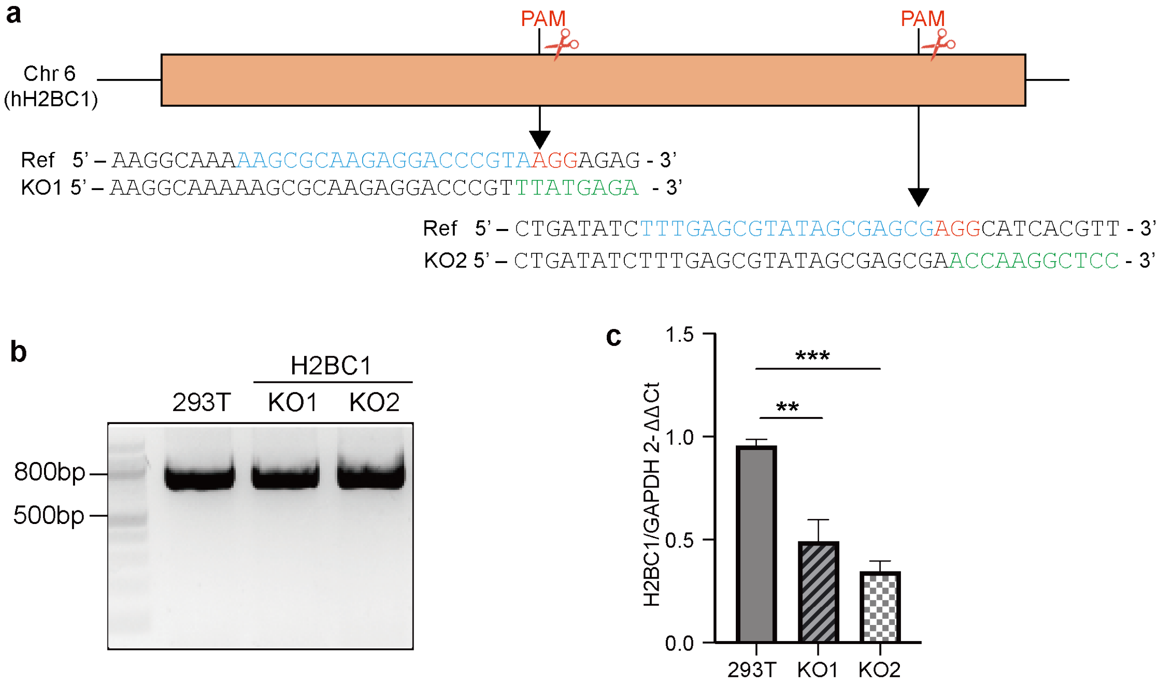


**Figure S6. Strategies for generating H2BC1 KO human 293T cell lines.**

**a.** Schematic diagram of guide RNA (gRNA) design targeting human H2BC1 gene and sequencing confirmation of generated stable clones. The sgRNA-targeting sequences are highlighted in blue, and PAM sequences are in red. The below schematic diagram shows the genotype of two correct targeted clones (KO1, KO2) and the mutated DNA sequences are shown in green.

**b.** PCR products of the genomic regions containing the targeting sequences were used for Sanger sequencing.

**c.** Relative expression level of H2BC1 in control and two 293T cell lines. The level of gene expression in control 293T cells was set as 1 (n = 3).

**Supplementary Figure 7**

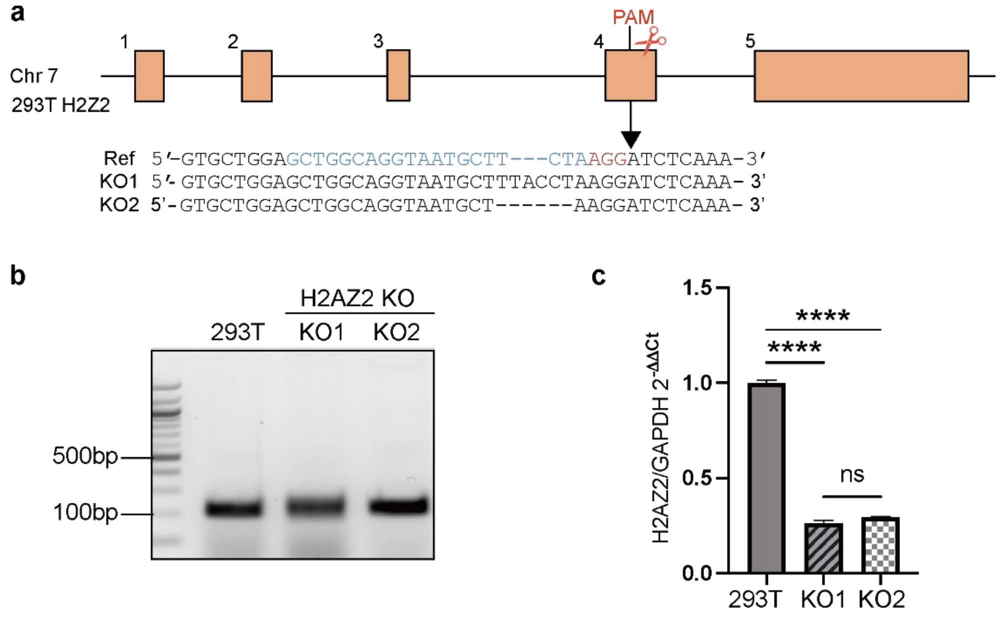


**Figure S7. Strategies for generating H2AZ.2 KO human 293T cell lines.**

- **a.** Schematic diagram of guide RNA design targeting H2AZ.2 and sequencing confirmation of generated stable clones. The sgRNA-targeting sequences are highlighted in blue, and PAM sequences are in red. The below schematic diagram shows the genotype of the correct targeted clones (#44 and #77).
- **b.** Genotyping analysis of H2AZ.2 KO 293T cells by PCR (n = 3).
- **c.** Relative expression level of H2AZ.2 in control and two 293T cell lines. The level of gene expression in control 293T cells is set as 1 (n = 3).

### Supplementary Figure 8

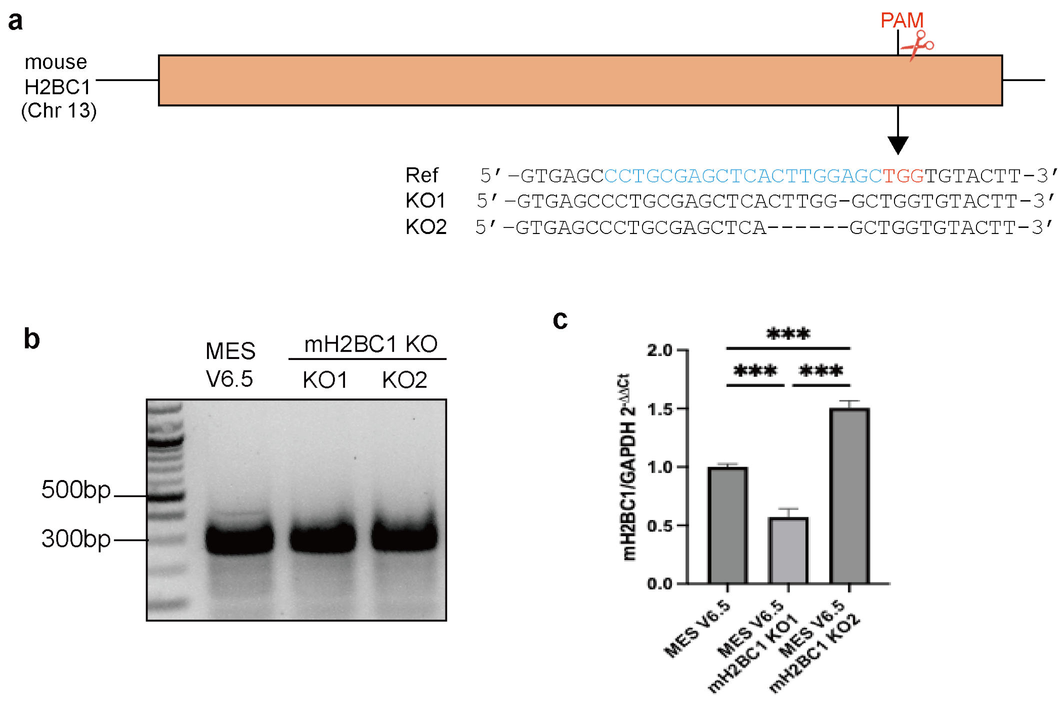


**Figure S8. Strategies for generating H2BC1 KO mouse ESC cell lines.**

- **a.** Schematic diagram of guide RNA design targeting H2BC1 and sequencing confirmation of generated stable clones. The sgRNA-targeting sequences were highlighted in blue, and PAM sequences are in red. The below schematic diagram shows the genotype of the correct targeted clones.
- **b.** Genotyping analysis of H2BC1 KO mouse ESC lines by PCR (n = 3).
- **c.** Relative expression level of H2BC1 in control and two ESC lines. The level of gene expression in control 293T cells is set as 1 (n = 3).

### Supplementary Figure 9

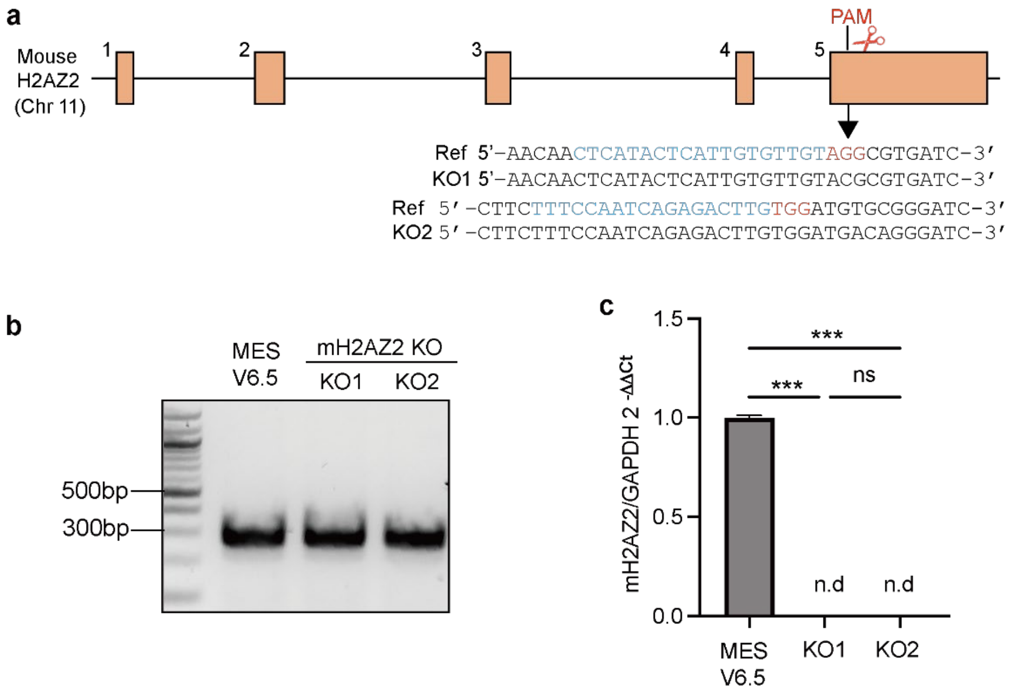


**Figure S9. Strategies for generating H2AZ.2 KO mouse ESC cell lines.**

**a.** Schematic diagram of guide RNA design targeting H2BC1 and sequencing confirmation of generated stable clones. The sgRNA-targeting sequences are highlighted in blue, and PAM sequences are in red. The below schematic diagram shows the genotype of the correct targeted clones.

**b.** Genotyping analysis of H2AZ.2 KO mouse ESC lines by PCR (n = 3).

**c.** Relative expression level of H2AZ.2 in control and two ESC lines. The level of gene expression in control mouse mESC is set as 1 (n = 3).

### Supplementary Figure 10

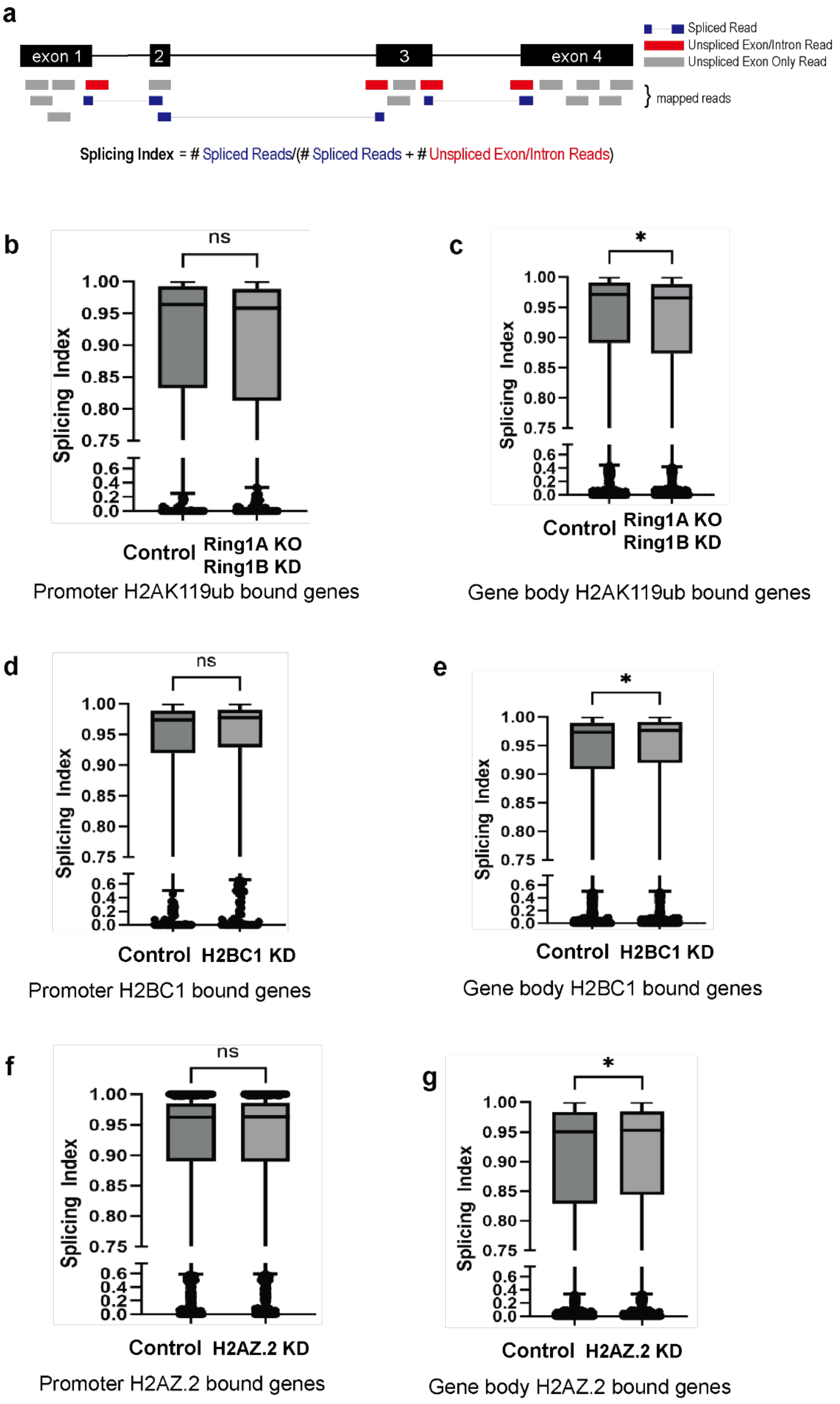

**Figure S10. Effects of Ring1AKO/Ring1BKD, H2BC1 KD, and H2AZ.2 KD on the gene-splicing efficiency in mESC.**

**a.** Diagram elucidating the splicing analyses. Splicing index is a measure of splicing competences, calculated by comparing the number of reads indicative of splicing to the total number of informative reads. The measure is bounded between 0 and 1, with 1 indicating a completely spliced transcript.

**b-g**. Boxplot for splicing index for genes bound by H2AK119ub (b, c), H2BC1 (d, e), and H2AZ.2 (f, g) at promoter region (b, d, f) and coding region (c, e, g). Note that Ring1AKO/Ring1BKD, H2BC1 KD and H2AZ.2 KD resulted in decreased splicing efficiency only when bound to the gene bodies of respective genes.

**Table S1. Cryo-EM Data Collection, Refinement, and Validation Statistics.**

| Description | Nucleosome*** |
|---|---|
| **EMDB** | *** |
| **PDB** | *** |
| **Microscope** | Krios |
| **Voltage (kV)** | 300 |
| **Camera** | K3 |
| **Magnification** | 36,000 |
| **Pixel Size (Å)** | 0.536 |
| **Spherical aberration (mm)** | 2.7 |
| **Total Electron Exposure (e-/A$^2$)** | 40 |
| **Defocus Range (μm)** | -0.9 ~ -2.5 |
| **Symmetry Imposed** | C1 |
| **Raw moives** | 5630 |
| **Initial Particles** | 4,887,804 |
| **Final Particles** | 1,288,371 |
| **Initial models** | 1AOI |
| **Map Resolution** (FSC threshold of 0.143) | 2.63 |
| **Map sharpening B-factor (A$^2$)** | 1.149 |
| **Chain** | 6 |
| **Non-hydrogen atoms** | 21187 |
| **Protein residues** | 766 |
| **Bonds (RMSD)** | |
| **Length (Å)** | 0.005 (0) |
| **Angles ($^o$)** | 0.730 (3) |
| **Molprobity score** | 1.60 |
| **Clashscore** | 5.58 |
| **Ramachandran plot (%)** | |
| **Outliers** | 0.13 |
| **Allowed** | 3.33 |
| **Favored (%)** | 96.53 |
| **Rama-Z (Ramachandran plot Z-score, RMSD)** | |
| **whole (N = 1884)** | 2.00 (0.30) |
| **helix (N = 73)** | 2.27 (0.22) |
| **sheet (N = 395)** | --- (---) |
| **loop (N = 1416)** | -0.93 (0.38) |
| **Rotamer outliers (%)** | 1.25 |
| **Cβ outliers (%)** | 0.00 |
| **Peptide plane (%)** | |
| **Cis proline/general** | 0.0/0.1 |
| **Twisted proline/general** | 0.0/0.0 |
| **CaBLAM outliers** | 1.23 |
| **ADP (B-factors)** | |
| **Iso/Aniso (#)** | 11663/0 |
| **min/max/mean** | |
| **Protein** | 0.00/66.49/18.01 |
